# DNA sequence sets the destination of spontaneous epimutations in plants

**DOI:** 10.64898/2026.10.21.760629

**Authors:** Ding Wang, Junru Jin, Jianbo Qiao, Leyi Wei, Qiang Liu

## Abstract

Plant genomes gain and lose cytosine methylation spontaneously, far faster than DNA mutates, and these epimutations are inherited. The genome is thought to set a steady state around which methylation fluctuates. Here we show that this steady state, the set point, is written in the DNA of each CG site: REED, a deep-learning model trained on one methylome, reads it. In pedigrees of seven *Arabidopsis thaliana* accessions, gains concentrated at high set points and losses at low ones, and the equilibrium implied by their rates matched the set point, tracking it within genes. Stress sped epimutation but barely shifted its destination. The set point predicted germline epimutations in rice and changes accumulated in the wild and in a 330-year-old tree. Its rules, cooperation between nearby CG sites and transcription-factor motifs that lower methylation, are shared across flowering plants, so a genome and one methylome reveal where epigenetic variation is heading.

## Introduction

Genomes carry heritable information in two layers: the DNA sequence and the chemical marks laid on top of it. In plants the second layer is restless. CG sites in *Arabidopsis thaliana* gain or lose methylation spontaneously at about 10^−4^ per site per generation, orders of magnitude above the rate of DNA point mutation, and the new states are transmitted through seed^1–4^. These epimutations recur at the same sites in independent lineages^5^, accumulate along the branches of long-lived trees^6^ and tick regularly enough to date plant lineages^7^. Whether this variation drifts freely or is pulled back towards a state written in the DNA decides how much heritable information it can carry beyond the sequence.

One view holds that the genome sets a steady state around which random epimutations make methylation fluctuate^8,9^. Epimutation rates imply a genome-wide equilibrium close to the average methylation of Col-0^3,10^, and a stochastic model of cooperative maintenance assigns each gene-body segment its own steady state^11^. CG density and repetitiveness predict which loci regain methylation after its loss in *met1* mutants^12^. For an individual cytosine, however, the steady state can at present be estimated only from the epigenome itself, such as methylation variability across hundreds of accessions^13^ or chromatin states^14^. Such estimates cannot tell whether a site’s steady state is carried by the chromatin or written in its DNA, and the data they need exist for almost no plant other than *A. thaliana*.

The steady state of a single site, which we call its set point, can be estimated in two independent ways. Mutation-accumulation lines, propagated for many generations by single-seed descent, give a time-lapse of the methylome: counting gains and losses at each site gives its equilibrium. A single methylome gives a snapshot. If gain and loss rates are set by the sequence, sites that share a sequence context flip like copies of one biased coin, and at any moment the fraction of them that is methylated is their set point; a sequence model trained on one methylome^15–19^, built on genomic language models pretrained across species^20–22^, should then learn it. Here we show that it does. We built REED (reading epigenetic equilibria from DNA), which reads 10 kb of DNA and predicts the methylation of every cytosine. In *A. thaliana* pedigrees, with or without stress, and in the rice germline, the equilibrium implied by spontaneous gains and losses matches REED’s prediction, and changes accumulated in the wild and in a 330-year-old poplar follow it. REED has learned two kinds of rule, cooperation between nearby CG sites and motifs whose binding proteins exclude methylation; both are shared across fourteen plants, and a model trained on one species predicts the direction of epimutation in another.

## Results

### REED reads a methylation set point from DNA

REED fine-tunes NTv3^21^, a DNA language model pretrained on many genomes, to predict the methylation level of every cytosine on both strands from 10 kb of sequence (Fig. 1a). The *A. thaliana* model was trained on chromosomes 1–3 of the consensus methylome of 605 accessions^23^, the least noisy labels; the models for all other species, and a second *A. thaliana* model used to repeat every epimutation test, were each trained on the methylome of a single plant.

**Fig. 1.**
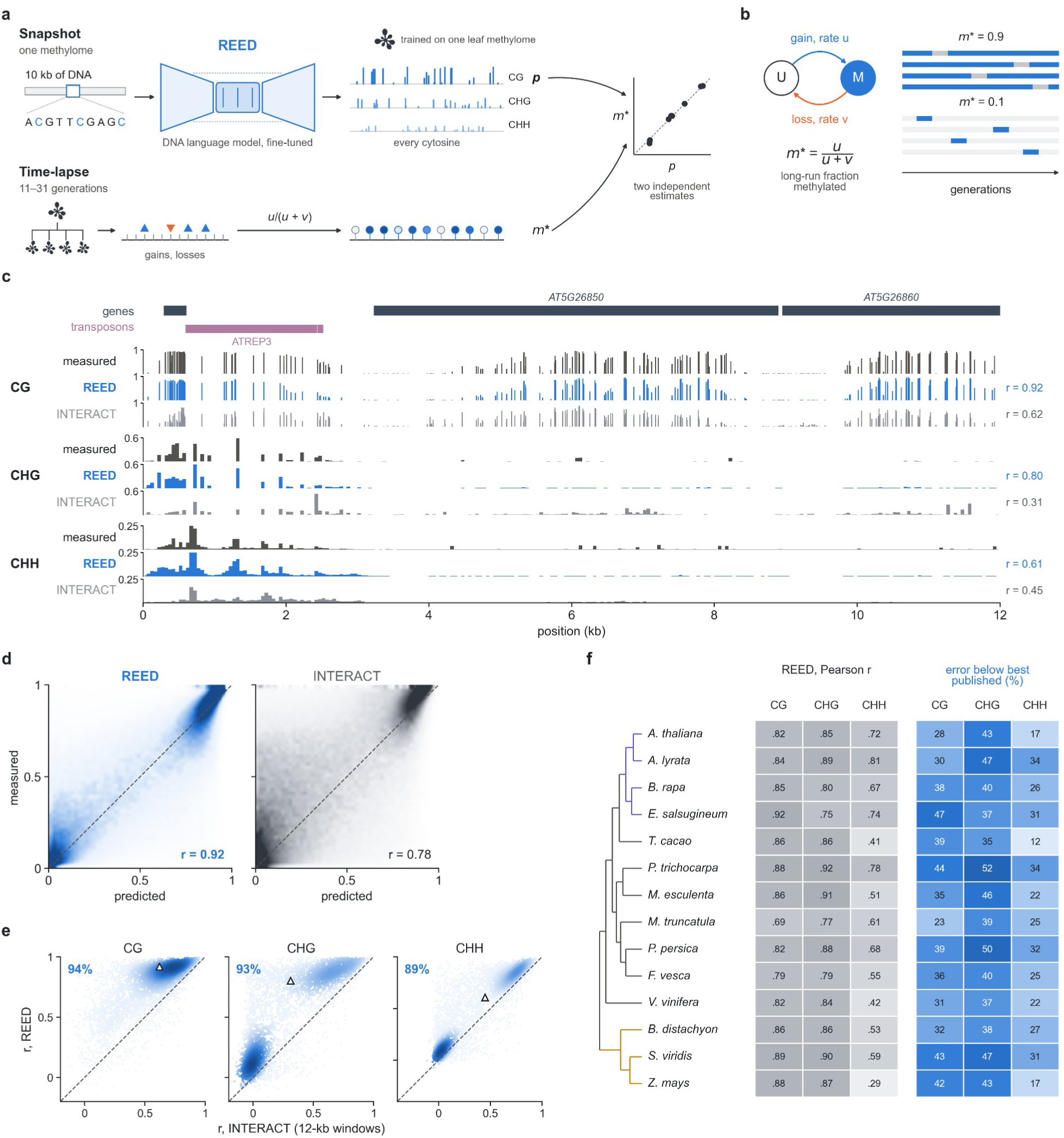
| REED reads a methylation set point from DNA. **a**, The framework. Snapshot: REED, a DNA language model fine-tuned on one leaf methylome, reads 10 kb of DNA and predicts the methylation of every cytosine; its prediction at each CG site is the set point *p*. Time-lapse: gains and losses counted in mutation-accumulation lines give each site’s equilibrium *m*\*. The two are independent estimates of the same quantity (one model per species; the *A. thaliana* model in **c**–**e** on the consensus of 605 accessions). **b**, A CG site switches between unmethylated (U) and methylated (M) states at rates *u* and *v*; its long-run methylated fraction is *m*\* = *u*/(*u* + *v*). Strips, four lineages each for a site with a high and a low set point (blue, methylated). **c**, Measured and predicted methylation over 12 kb of held-out chromosome 5 (every CG site; 50-bp means for CHG and CHH) for REED and INTERACT; right, correlations in this window. **d**, Measured against predicted CG methylation for all 1.2 million CG sites of chromosome 5; each row shows where predictions fall for sites at that measured level; numbers, Pearson *r* over all CG sites. **e**, Pearson *r* of REED against that of INTERACT in each 12-kb window of chromosome 5, by context (points coloured by density); numbers, percentage of windows in which REED is more accurate (above the diagonal); triangles, the window in **c**. **f**, Fourteen species: left, Pearson *r* of the REED species model on held-out regions (for *A. thaliana*, the model trained on a single plant); right, how much lower REED’s mean squared error is than that of the best of three published methods retrained on each species (%); rows follow the phylogeny (Brassicaceae violet, grasses gold).

A CG site switches between unmethylated and methylated states, gaining methylation at rate *u* and losing it at rate *v*; left long enough, it spends a fraction *m*\* = *u*/(*u* + *v*) of the time methylated (Fig. 1b), its set point. Pedigrees give *m*\* by counting gains and losses; REED gives an estimate of it, *p*, from sequence (Fig. 1a), and from here on we call *p* the set point.

On held-out chromosome 5, REED reproduced methylation down to individual CG sites (Pearson *r* over the chromosome, 0.92, 0.92 and 0.83 for CG, CHG and CHH; one window in Fig. 1c, all CG sites in Fig. 1d), was more accurate than INTERACT^16^, a published deep-learning model, in 89–94% of 12-kb windows (Fig. 1e) and had 38–70% lower error than three published methods retrained on the same data^15–17^ (Supplementary Fig. 1). REED models for fourteen species, the thirteen beyond *A. thaliana* each trained on one public leaf methylome^24–26^ and including three grasses and the crops maize, cassava and strawberry, were more accurate than the same three methods retrained on each species in all 42 species–context combinations, with the largest margins where CHH methylation is sparse (Fig. 1f and Supplementary Fig. 2).

### Spontaneous epimutations move towards the set point

We compared *p* with spontaneous CG epimutations in two independent Col-0 pedigrees: the Shaw population, nine lines propagated for 31 generations and sequenced by two laboratories^1–3,27^, and MA3, two lineages followed for 11 generations from a sequenced ancestor^28^ (Fig. 2a). All tests used chromosomes 4 and 5, unseen in training, and were specified in advance. At a single gene (Fig. 2b), AT5G51340, unmethylated CG sites that gained methylation in one of the nine lines had a mean set point of 0.57, against 0.17 for those that did not.

**Fig. 2.**
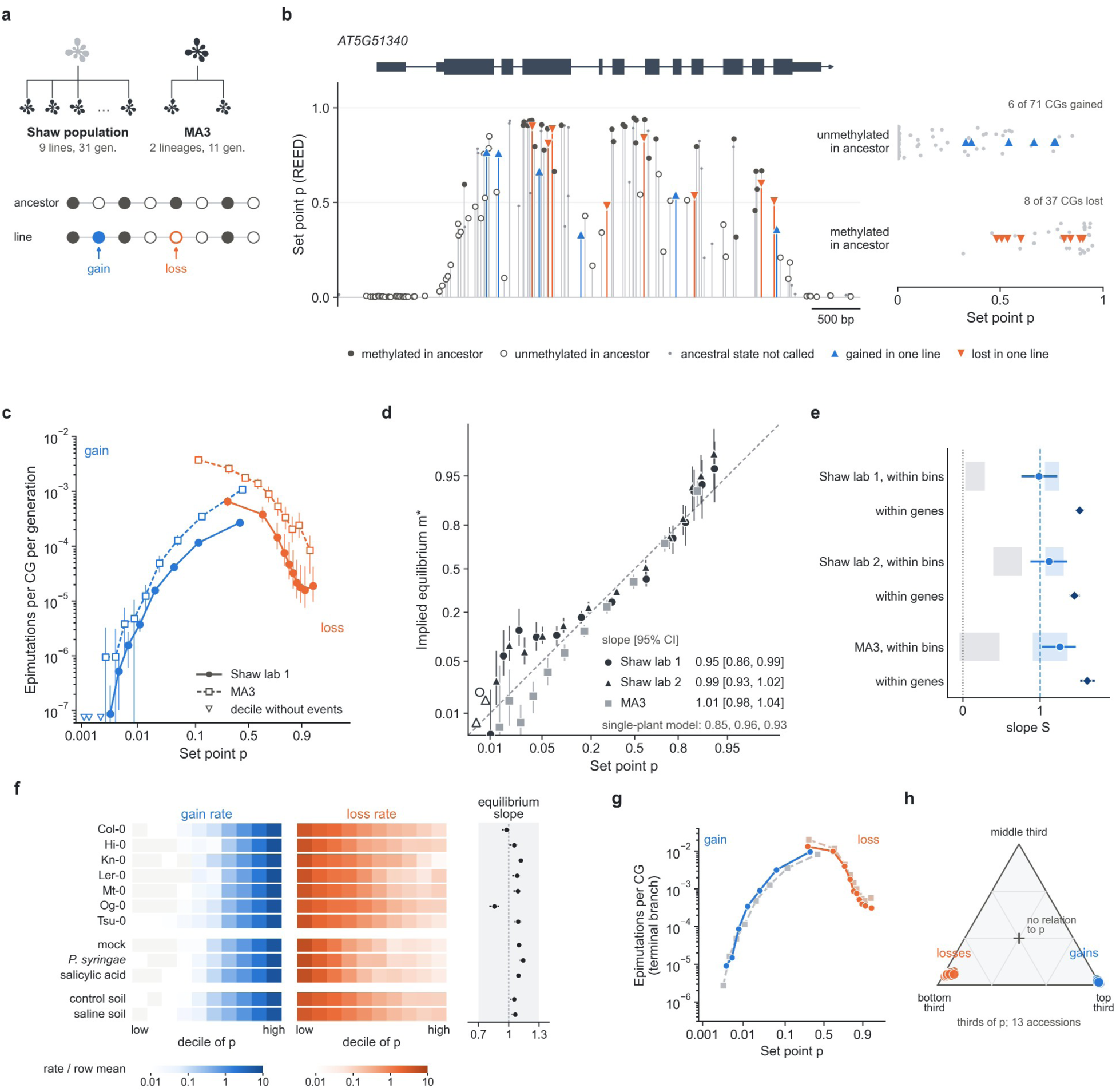
| Spontaneous epimutations move towards the set point. **a**, The two mutation-accumulation designs: the Shaw population (nine lines, 31 generations; ancestral states inferred from the other lines) and MA3 (two lineages, 11 generations; ancestor sequenced). **b**, AT5G51340 (held-out chromosome 5): each CG at the height of its set point *p*, with its ancestral state and the gains (blue) and losses (orange) in nine Shaw-population lines; right, the gene’s ancestrally unmethylated and methylated CGs placed by *p*. **c**, Gain rate of ancestrally unmethylated CGs and loss rate of ancestrally methylated CGs per generation against *p* (deciles; logit axis) for the Shaw population (laboratory 1; filled) and MA3 (open). **d**, Equilibrium implied by epimutation rates, *m*\* = *u*/(*u* + *v*), against *p* in 20 bins for the Shaw population sequenced by two laboratories and for MA3 (chromosome 5; logit axes); dashed line, identity; open symbols, bins whose bootstrap upper limit is undefined; below the key, slopes for the model trained on a single plant. **e**, Slope *S* of the rate-implied equilibrium on *p* among sites of the same bin of *p* (circles) or gene (diamonds); dashed line, *S* = 1, expected if *p* is each site’s set point; dotted line, *S* = 0; bands, ranges over simulations in which sites of a bin share one set point (grey) or in which *p* is each site’s set point (blue). **f**, Gain (blue) and loss (orange) rates by decile of *p*, relative to the mean rate of each row (log scale; light grey, deciles without events), in mutation-accumulation lines of Col-0, six further accessions and stress treatments with their controls (chromosomes 4 and 5); right, equilibrium slope of each row (shaded, 0.7–1.3, the range counted as agreement). **g**, Private gains and losses per terminal branch in 13 accessions of the North American haplogroup-1 lineage against *p* (coloured; grey dashed, Shaw population, per line over its 31 generations). **h**, Each accession’s private gains (blue) and losses (orange) as points in a ternary plot of the shares in the bottom, middle and top thirds of *p*; cross, equal shares, expected if epimutation ignored *p*. Error bars, 95% confidence intervals (1-Mb block bootstrap).

Across the genome, the gain rate of unmethylated sites rose more than a thousandfold with *p* and the loss rate of methylated sites fell 40-fold (Fig. 2c). Unmethylated sites in the top third of *p* gained methylation about 2,500 times more often than those in the bottom third, and methylated sites in the top third lost it 21 times less often. Among sites with the same neighbouring methylation, each standard deviation of *p* raised the odds of gain by 70% and lowered the odds of loss by 42%. The result was the same in both laboratories’ data, in MA3 after accounting for the founder’s own CHG and CHH methylation, and among CG sites of the same gene.

The most direct test is the equilibrium. Across 20 bins of *p*, the equilibrium implied by the counted rates matched *p* in value, not only in rank, from 0.01 to 0.95, with slopes of 0.95 and 0.99 in the two laboratories’ data and 1.01 in MA3 (Fig. 2d); the single-plant model gave 0.85, 0.96 and 0.93. The match holds site by site. Within narrow bins of *p*, sites with slightly higher *p* gained methylation more often and lost it less often, so that the slope *S* of the rate-implied equilibrium on *p* within bins was 0.99–1.26, and 1.45–1.61 within genes (Fig. 2e); simulations without site-level set points gave values far below these (Supplementary Fig. 3).

The set point does not belong to one genotype or one environment. In mutation-accumulation lines of six further accessions^7^, gains rose and losses fell with *p* in every accession (each standard deviation of *p*: odds of gain +80%, of loss −55%), and the equilibrium again matched the set point (pooled slope 0.98; Fig. 2f). Salicylic acid and soil salinity^7,29^ accelerated gains and losses by 21–40% at the same set point (infection with *Pseudomonas syringae* did not), whereas the effect of *p* and the equilibrium slope (1.05–1.14) were unchanged; because gains and losses accelerated nearly in proportion, the equilibrium moved by at most 0.06 under either treatment, within a preregistered equivalence margin of 0.1.

Epimutations accumulated in nature followed the same set points. The haplogroup-1 lineage of *A. thaliana* spread across North America within the past few centuries^5^. In 13 of its accessions, private gains were about 4,600 times more frequent in the top than in the bottom third of *p* and private losses 23 times less frequent, and the pattern held in every accession (Fig. 2g,h).

Among predictors that need no population data, REED was the most accurate: it predicted these epimutations better than published sequence models, gradient-boosting models built on sequence and annotation features and the cooperative-maintenance model of Briffa et al.^11^, and it added information to chromatin states^14^; the model trained on a single plant also outperformed the published models, and REED’s advantage held at sites with no detectable similarity to the training chromosomes (Supplementary Figs. 4 and 5). Only predictors read from the population itself, methylation frequency across the 605 accessions and its variability^13^, ranked sites as well or better; such panels exist for almost no other plant.

### The set point can be read across plants

Across every system in which spontaneous change could be measured, from *A. thaliana* pedigrees and wild plants to the germline of a crop and the soma of a tree, the set point predicted its direction with similar effect sizes: each standard deviation of *p* raised the odds of gain by 59–124% and lowered the odds of loss by 42–58% (Fig. 3a).

**Fig. 3.**
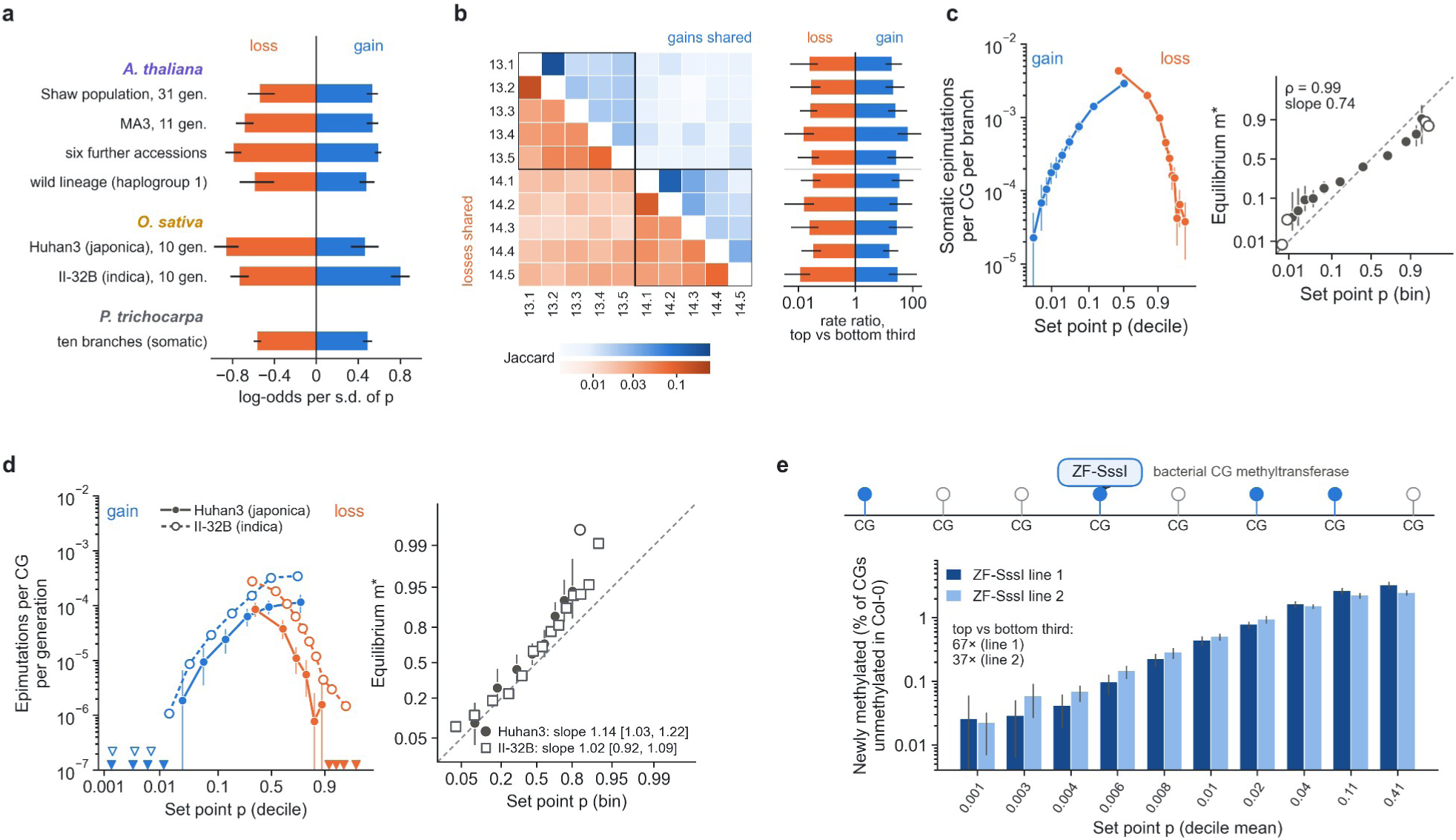
| The set point can be read across plants. **a**, Effect of the set point on spontaneous gains (blue) and losses (orange) in every system where change was measured, as log-odds per standard deviation of *p* controlling for neighbouring methylation, on held-out chromosomes. **b**, Somatic epimutations among ten branches of a wild *P. trichocarpa* tree about 330 years old; the poplar model was trained on a different tree. Left, epimutations shared by two branches (Jaccard index; gains above and losses below the diagonal; branches labelled by trunk and branch; boxes, the two trunks); right, each branch’s gain (blue) and loss (orange) rate at sites in the top versus the bottom third of the poplar set point. **c**, Single-branch somatic gain and loss rates by decile of *p* and the implied equilibrium after correction for false-positive calls, on test chromosomes 17–19. **d**, Germline epimutations in two rice lineages propagated by single-seed descent for ten generations under repeated drought, Huhan3 (japonica; filled, solid) and II-32B (indica; open, dashed): gain and loss rates per generation by decile of the rice set point (triangles, deciles without events) and the implied equilibrium (Huhan3 circles, II-32B squares), on chromosomes 11 and 12, held out from the rice model trained on one Nipponbare leaf methylome. **e**, Percentage of CG sites unmethylated in all six Col-0 controls that became methylated in two ZF-SssI lines, by decile of *p* (y axis, log scale); top, schematic. Grey dashed lines, identity; open circles, bins whose bootstrap upper limit is undefined (in **d**, for Huhan3). Error bars, 95% confidence intervals (1-Mb block bootstrap).

In a wild *Populus trichocarpa* tree about 330 years old, ten branches from two trunks record somatic epimutations accumulated over decades to centuries^6^. A poplar model trained on a different tree^24^ and run on the old tree’s own assembly predicted them on chromosomes held out from training. Unmethylated sites in the top third of *p* gained methylation 22 times more often than those in the bottom third, methylated sites in the top third lost it 39 times less often, the relationship held in every branch (Fig. 3b), and the implied equilibrium rose monotonically with *p* (Spearman ρ = 0.99; slope 0.74 when all other branches had to agree on the ancestral state, and 0.96 genome-wide when one or two could disagree; Fig. 3c and Supplementary Fig. 6).

A crop germline gave the same answer. The rice (*Oryza sativa*) cultivar Huhan3 was propagated by single-seed descent for ten generations under repeated drought^30^. We predicted set points with a fifteenth REED model, trained for rice on one leaf methylome of another cultivar, Nipponbare^31^, and tested on chromosomes held out from training. Not one of 330 gains fixed in the lineage occurred among unmethylated sites in the bottom third of *p*, and not one of 180 losses among methylated sites in the top third; each standard deviation of *p* raised the odds of gain by 59% and lowered the odds of loss by 58%, and the implied equilibrium matched the set point (slope 1.14; Fig. 3d). The same held for normally watered siblings alone and for a second lineage, of the indica cultivar II-32B (odds of gain +124% and of loss −52% per standard deviation; equilibrium slope 1.02; Fig. 3d).

The set point also predicted where a foreign enzyme succeeds. In two *A. thaliana* lines expressing the bacterial CG methyltransferase SssI fused to a zinc finger^32^, CG sites in the top third of *p* became methylated 67 and 37 times more often than those in the bottom third (AUC 0.82 and 0.79; Fig. 3e). The set point thus reports how readily a site accepts methylation, whether spontaneously or from a bacterial enzyme.

### What sets the set point

Two kinds of rule in the DNA determine the set point. In silico, removing the other CG sites within 100 bp of a gene-body CG with a high set point lowered its predicted methylation by 0.39, and adding a single CG 3–15 bp from a site with an intermediate set point raised its prediction by 0.17; the effect decayed within 60 bp and was nearly absent at sites with a low set point (Fig. 4a). REED has learned that CG sites stabilize one another over short distances where methylation is otherwise permitted, as expected from cooperative maintenance by MET1^11^. Natural variation confirmed this rule. Among 586 accessions^23,33^, a polymorphism that created a CG 3–14 bp from a methylable site (*p* ≥ 0.3) raised its methylation by 0.12 and one that destroyed a CG lowered it by 0.16, with no effect beyond 60 bp. Across 294,969 variant–site pairs the observed effects followed REED’s predictions (*r* = 0.54, slope 0.82; Fig. 4b) more closely than those of published methods, and the effects held among carriers and non-carriers with identical local haplotypes (Supplementary Figs. 7–9).

**Fig. 4.**
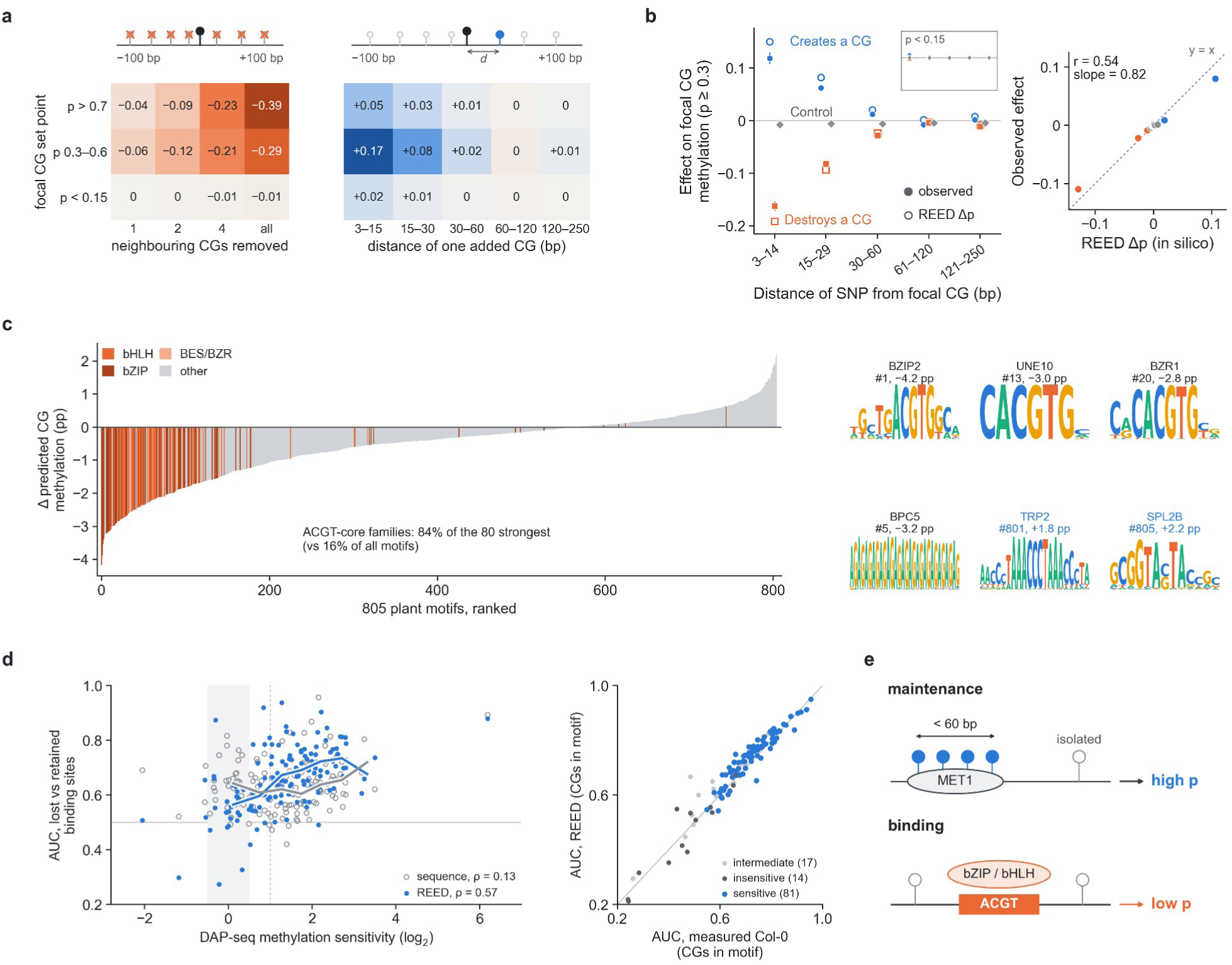
| What sets the set point. **a**, In silico CG cooperativity at gene-body CGs grouped by set point (rows): change in predicted methylation after removing neighbouring CGs within ±100 bp (left) or adding one CG at distance *d* (right); mean of 200 focal CGs per group. **b**, Natural variants among 586 accessions that create (blue) or destroy (orange) a CG near a methylable gene-body CG change its methylation as REED predicts (open symbols); grey, control variants; inset, focal CGs with low set points on the same scale. Right, observed against predicted effects in 20 bins of 294,969 variant–site pairs; *r* and slope over all pairs. **c**, Change in predicted CG methylation after inserting each of 805 plant motifs into moderately methylated *A. thaliana* sequences (mean of two models), ranked; ACGT-core families in shades of orange; right, logos of named examples with rank and effect: the strongest motif of each ACGT-core family, the strongest lowering motif without CG in its consensus and the two strongest raising motifs. **d**, Left, for each *A. thaliana* factor with DAP-seq data, AUC for separating sites bound on amplified DNA but lost on native genomic DNA from those retained, using REED’s predicted CG methylation within ±100 bp (blue) or a cross-validated sequence-composition classifier (grey, open), against the factor’s methylation sensitivity; lines, running medians; shaded, insensitive factors; dotted line, twofold reduction. Right, the same comparison at the CG sites inside each factor’s motif hit, REED against the measured Col-0 methylome (112 factors with CG-containing motif hits). **e**, Schematic of the two routes from sequence to set point suggested by **a**–**d**: cooperative maintenance across neighbouring CG sites raises the set point, and transcription factors bound at ACGT-core motifs exclude methylation and lower it. Error bars, 95% confidence intervals (1-Mb block bootstrap).

These rules act within a background that REED reads from kilobases of DNA. When the sequence beyond 250 bp of a gene-body CG site was swapped between genes with gene-body methylation (gbM) and unmethylated genes, methylated sites lost 96% of their higher set point, whereas unmethylated sites gained only 26%; shuffling the distant flanks of methylated sites while keeping every dinucleotide frequency removed 55% (Supplementary Fig. 10). The methylable regions that the cooperative-maintenance model takes from chromatin data^11^ are thus written in the arrangement of gene sequence.

Sequence motifs add a second set of rules. Inserted into moderately methylated sequences, motifs with an ACGT core, bound by bHLH, bZIP and BES/BZR factors, lowered predicted CG methylation by 2–4 percentage points; these families made up 84% of the 80 motifs that lowered methylation most, against 16% of all 805 plant motifs^34^ (Fig. 4c and Supplementary Fig. 11). Factor binding and methylation exclude each other: many of these factors cannot bind methylated DNA^35^, and REED’s set point marks the sites where methylation wins. For 90 factors whose DAP-seq binding is at least halved by CG methylation, REED’s prediction separated binding sites lost on native genomic DNA from those retained (median AUC 0.70, against 0.67 for a measured Col-0 methylome; above 0.5 for 89 of 90 factors), whereas it carried little information for factors insensitive to methylation (Fig. 4d).

### A methylation grammar shared across angiosperms

Each species model learned its grammar from that species’ own methylome, so the grammars of species separated by up to 160 million years can be compared directly (Fig. 5a). As families, bZIP motifs lowered predicted CG methylation in every species except strawberry, and bHLH motifs in every species except maize; their methylomes show the weakest ACGT footprints of the fourteen (Supplementary Fig. 12). Of the 805 motifs, 288 changed predicted methylation in the same direction in at least 12 species, and half of these had a footprint in the same direction around native motif instances in at least eight species’ measured methylomes, against a quarter of other motifs (Fig. 5b). Besides the ACGT core, MADS, NAC, TCP, SBP and ERF motifs lowered methylation, and telomere-repeat, GATA and AIL motifs raised it.

**Fig. 5.**
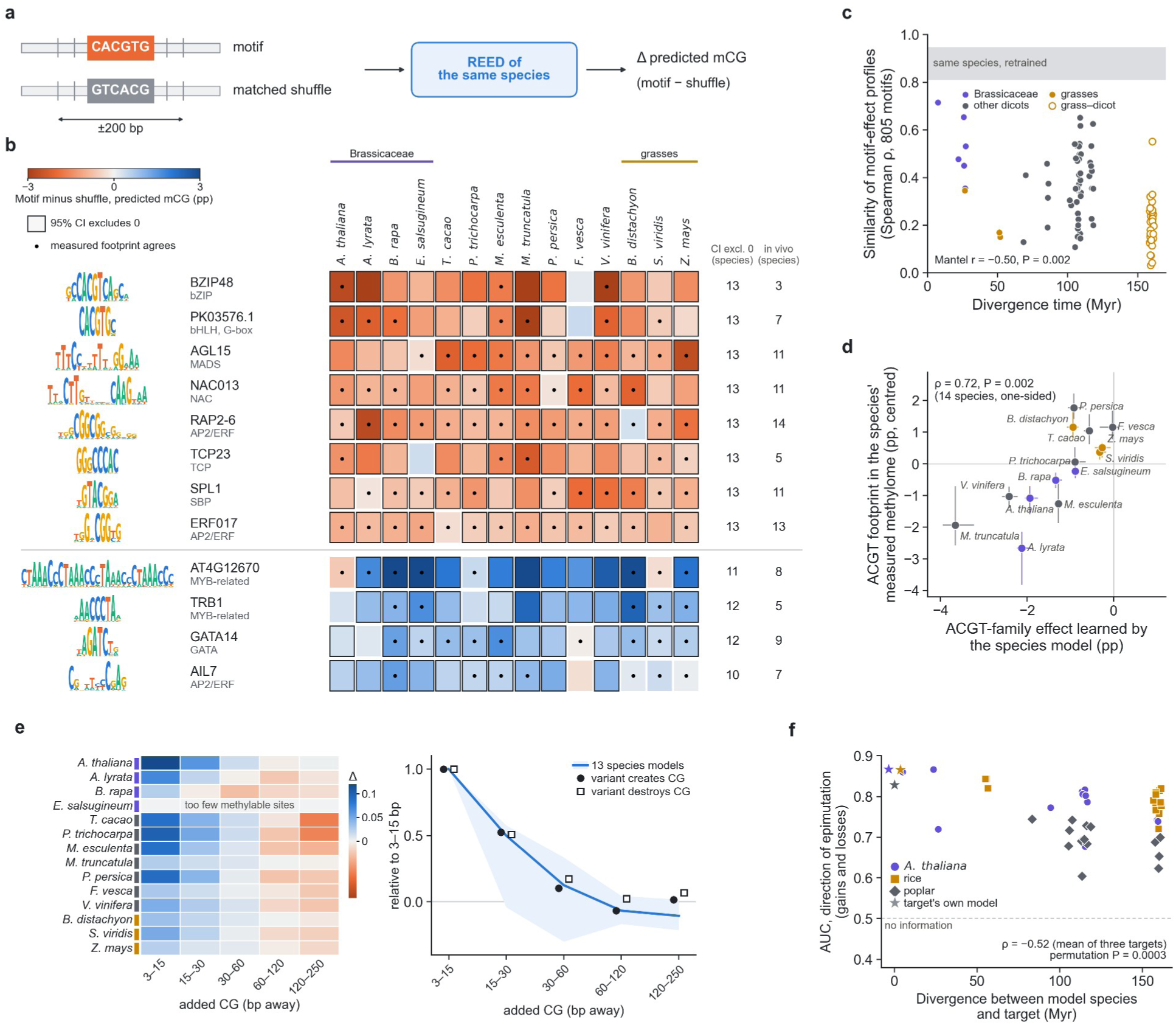
| A methylation grammar shared across angiosperms. **a**, Design: each of 805 JASPAR plant motifs and a matched shuffle are inserted into 10-kb backgrounds from each species’ own genome and scored by that species’ REED model as the change in predicted CG methylation within ±200 bp. **b**, Representative motifs with conserved effects across fourteen species, one per cluster of related motifs: logo, name and family, effect in each species (200 moderately methylated backgrounds per species; frame, 95% bootstrap interval excluding zero; dot, measured footprint around native instances in the same direction), and the numbers of species with intervals excluding zero and with agreeing footprints. **c**, Similarity of motif-effect profiles between species pairs against divergence time (violet, within Brassicaceae; dark grey, other dicot pairs; gold, within grasses; open gold, grass–dicot pairs); grey band, similarity between independently retrained models of the same species; one-sided Mantel test over 14 species. **d**, Strength of the ACGT-family effect learned by each species model against the ACGT footprint in its measured methylome, centred on the median of all motifs (colours as in **c**). **e**, CG cooperativity read by fourteen species models on the same DNA. Left, change in predicted methylation of methylable *A. thaliana* gene-body CG sites (*p* ≥ 0.3 under each model) after adding one CG at the indicated distance, minus a matched substitution that creates no CG (rows, species in phylogenetic order; *E. salsugineum*, too few methylable sites to test); right, the same effects scaled to each model’s 3–15 bp effect (band, range over 13 models; line, median), with natural variants that create (filled) or destroy (open) a CG among 586 *A. thaliana* accessions scaled the same way. **f**, Accuracy of borrowed set points in predicting the direction of epimutation (mean of gain AUC and one minus loss AUC) in *A. thaliana* (violet), rice (gold) and poplar (dark grey) against divergence time between model species and target; stars, the target’s own model. Error bars in **d**, 95% bootstrap intervals.

The similarity of motif-effect profiles between species declined with divergence time, mainly across the split between monocots and dicots (Mantel *r* = −0.50, *P* = 0.002; Fig. 5c), and the strength of each model’s ACGT effect followed the depth of the ACGT footprint in its own methylome (Spearman ρ = 0.72, *P* = 0.002; Fig. 5d). In the *A. thaliana* pedigrees, CG sites near conserved ACGT-core motifs gained methylation less often than others with the same set point (rate ratio 0.83, 0.73–0.95).

CG cooperativity is shared as well. Given the same 300 *A. thaliana* gene-body CG sites with the same added or removed neighbouring CGs, 12 of the 14 species models, including all three grasses, raised the predicted methylation of a methylable site when a CG was added 3–15 bp away and lowered it when neighbouring CGs were removed. The strength of the rule varied fivefold between species, but its range did not: in 13 models the effect fell to a median of half at 15–30 bp and vanished beyond 60 bp, the decay measured for natural variants among *A. thaliana* accessions (Fig. 5e). In each species’ own methylome, the measured short-range footprint ranked the species as their models did (Spearman ρ = 0.70, *P* = 0.004; Supplementary Fig. 13).

Because the grammar is shared, a model trained on one species predicts where epimutations go in another. Scoring the *A. thaliana*, rice and poplar epimutations with each of the other species’ models, the borrowed set point predicted gains and losses in the expected direction in 35 of 42 combinations, with accuracy declining with divergence time (Spearman ρ = −0.52, permutation *P* = 0.0003; Fig. 5f). Own-species models remained the most accurate.

## Discussion

Plants change their methylation spontaneously and pass the changes on, which has made them the strongest case for epigenetic variation as a second layer of inheritance. Our results tie that layer closely to the first. At CG sites, spontaneous epimutation behaves like a rarely flipped, biased coin, and the bias is written in the surrounding DNA. A model trained on leaf methylation predicts the direction and long-run outcome of germline change in two plant species, with or without stress, and the direction of change in nature, in an old tree and at sites targeted by a bacterial enzyme. The view that genotype sets a steady state around which methylation fluctuates^3,8,9,11^ thereby extends to single sites and becomes readable from sequence. The principle need not be confined to plants: in mammals, DNA fragments inserted at a common locus adopt methylation states dictated by their own sequence^36^, and individual CpG sites flip between states as somatic lineages grow^37^.

The rules REED has learned point to two physical routes from sequence to set point: the maintenance machinery, under which a CG site is stabilized by methylable neighbours within a few tens of base pairs^11^, and the proteins that bind DNA and exclude methylation from their sites^35^ (Fig. 4e). Models trained independently for fourteen species converge on the same core grammar, from the range of CG cooperativity to the motifs that lower methylation, with a strength that each species tunes. The set point is defined for the methylation enzymes a genome carries: variants at trans-acting loci shift methylation genome-wide^10,38^, and REED models trained on a species that has lost CMT3 predict methylation resembling that of a *cmt3* mutant, and the CMT2 rule each model learns follows its species’ methylome (Supplementary Fig. 14).

Knowing where a site is heading tells us whether a new epiallele will last. A deviation from the set point decays at rate *u* + *v*, over 165–550 generations in MA3 (median 289), so an epiallele behaves like a stable allele over a few generations, yet one that sits against its set point is expected to revert over centuries. The set point likewise tells an epigenome engineer which targets readily accept new methylation^32,39^, and it gives a per-site expectation under epimutation alone against which the effects of selection and environment become measurable. Because REED needs only a genome and one methylome, set points can now be read in crops and trees that have neither mutation-accumulation lines nor population panels.

## Methods

### Methylomes and genomes

The *A. thaliana* training labels were the per-cytosine consensus methylation of 605 leaf methylomes from the 1001 Epigenomes collection^23^ on the TAIR10 assembly, restricted to cytosines covered in at least 20 accessions. Each of the other species models was trained on a single public leaf methylome^24–26,31^, as was a second *A. thaliana* model (a Col-0 plant of the third generation of the Shaw population^2,25^). Where the assembly used in the original study differed from the NCBI release, reads were realigned with bwa-meth^40^ and methylation was extracted with MethylDackel. Chromosome splits were fixed from assembly structure before methylation values were read; in *A. thaliana*, chromosomes 1–3 were used for training, chromosome 4 for model selection and chromosome 5 for testing only, and in rice (Nipponbare mature leaf, first 150 million read pairs, aligned to IRGSP-1.0^41^), chromosomes 1–8 for training, 9–10 for model selection and 11 –12 for testing.

### REED and baselines

REED fine-tunes the publicly released NTv3-100M post-trained model^21^, a U-Net-like DNA language model with single-nucleotide resolution that was pretrained by masked-nucleotide prediction on many animal and plant genomes and then post-trained on functional-genomic tracks. A per-base head predicts the methylation of every cytosine on both strands from 10 kb of sequence. The loss was the mean squared error over the central 4,096 bp, with CG, CHG and CHH weighted 1:1:2; for models trained on a single methylome, each cytosine with at least five reads was weighted by its read coverage, capped at 20. After the head had been trained with the backbone frozen, all parameters were fine-tuned (8,000 steps, learning rate 3 × 10^−5^, batch size 32) in full-precision arithmetic, and checkpoints were selected on the validation chromosome. None of the post-training tracks is a methylation assay, maize and rice are the only other species of this study among the post-training species, and a REED model built on the backbone before post-training reproduced the epimutation results (gain AUC 0.886 against 0.896; equilibrium slope 0.91 against 0.95).

CpGenie^15^, INTERACT^16^ and the DNA-only configuration of EvoPlantMeth^17^ were retrained on the same labels and splits as REED, first on the *A. thaliana* consensus and then on each species; CpGenie predicts CG only. The annotation baseline was a gradient-boosting model^42^ on local sequence composition and gene and transposon annotation. A second REED seed and a variant with the 650M-parameter backbone are shown in Supplementary Figs. 1 and 4. Performance was measured as Pearson *r* and mean squared error over all cytosines with coverage ≥ 5 on held-out chromosomes. Examples shown in figures were chosen by rules fixed before plotting: the window in Fig. 1c had to contain genes and transposons and variable methylation, with REED’s advantage over INTERACT at the 85th percentile of all windows; the gene in Fig. 2b was, among 166 protein-coding genes of 1.5–6 kb with at least three gains and three losses, the one with the most events among those whose within-gene AUCs for gain and loss fell in the interquartile range and whose event count stayed below the 90th percentile.

### Epimutations in mutation-accumulation lines

Shaw-population methylomes^1,2^ were taken from the uniformly reprocessed data of ref.^3^ and MA3 methylomes from ref.^28^; analyses used chromosomes 4 and 5. A CG site was called methylated at ≥ 0.7 and unmethylated at ≤ 0.1. In the Shaw population, the ancestral state for each line was the concordant state of the other lines, and an epimutation required every callable replicate at generation 31 to show the opposite state; 99.0% of epimutations called in one laboratory’s data reappeared in the other’s. In MA3, the ancestral state was that of the sequenced founder, and an epimutation had to be present at both generations 8 and 11.

Gain rates *u* and loss rates *v* were estimated in bins of *p* as events per ancestrally unmethylated or methylated site per generation, and *m*\* = *u*/(*u* + *v*) was regressed on the bin mean of *p* with inverse-variance weights. Effects of *p* beyond neighbouring methylation (±100 bp) and within genes were estimated by logistic regression and are reported as changes in odds per standard deviation. For the site-level slope *S*, gains and losses were each regressed on logit *p* with fixed effects for bins or genes, so that *S* = β_gain_ − β_loss_ estimates the slope of logit *m*\* on logit *p* within strata; *S* above 1 within genes means that set points differ between sites of one gene even more steeply than REED predicts. Simulations placed two-state processes on the real sites, with the real coverage and line structure and rates calibrated to the observed event counts, and passed them through the same calling code. Predictors from the literature were evaluated on the same epimutations: chromatin states (36 states of the Plant Chromatin State Database) assigned to each CG site, with state-specific gain and loss rates estimated on chromosomes 1–3^14^; steady states of the cooperative-maintenance model, taken from the published segment-level predictions^11^; and neighbouring methylation combined with variability across the 605 accessions in a logistic model^13^. The relaxation time of a site is 1/(*u* + *v*) generations.

Mutation-accumulation lines of six further accessions and of Col-0 under biotic stress^7^ or soil salinity^29^ were analysed with the rules used for MA3 after removing CG sites within 50 bp of a sequence difference from TAIR10; *p* was taken from the reference prediction, and predictions on each accession’s own sequence gave the same results. For each stress treatment and its control, gain and loss rates were modelled within bins of *p* with a common stress factor or with separate factors for gains and losses; the shift of the equilibrium, Δ = log(*u*_stress_/*u*_control_) − log(*v*_stress_/*v*_control_), equals the change in logit *m*\* and was tested for equivalence against a preregistered margin of log 1.5 (a change of *m*\* of at most 0.1). In MA3, the founder’s mean CHG and CHH methylation within 100 bp was added to the logistic models as a covariate.

### Epimutations in nature, in rice, in poplar and after ZF-SssI

The same calling rules and code were applied to the natural lineage, the rice pedigrees and, with adaptations for single samples, to poplar. Bisulfite reads of 13 haplogroup-1 accessions^5^ were aligned to TAIR10, CG sites overlapping any sequence variant of these accessions^33^ were removed, and each accession’s private epimutations were called against the concordant state of the other twelve; 99.5–100% of the epimutations called in one library reappeared in a second library of the same accession. For rice, the 16 Huhan3 methylomes of ref.^30^ (founder generation, generations 10 and 11, and a generation grown from generation-10 seed without drought; two drought-treated and two normally watered siblings each) were realigned to IRGSP-1.0 with the same pipeline. The ancestral state of a CG site required agreement among the four founder siblings (at least three callable) and their pooled reads; an epimutation required the pooled calls of all three descendant groups to oppose it, with no callable descendant showing the ancestral state, so that changes arising after generation 9, which still segregate among siblings, were excluded. Rates assume ten generations. *p* was predicted on the Nipponbare reference sequence; no epimutation fell on a CG disrupted by a Huhan3 sequence variant detected from the bisulfite reads. The 16 methylomes of the indica lineage II-32B from the same study were processed identically with the same model; because indica differs more from the reference, CG sites disrupted by II-32B variants detected from its bisulfite reads (5.4% of CG dinucleotides on chromosomes 11 and 12) were masked in the primary analysis, and results without masking were the same. A sibling-based noise control showed more residual segregation in II-32B than in Huhan3; the direction and equilibrium results held with stricter descendant and ancestral rules. For the poplar tree^6^, the poplar model was run on the tree’s own assembly; each branch was compared with the concordant state of the other nine, a somatic epimutation required a change of at least 0.25, consistent on both strands, at depth ≥ 20, and the implied equilibrium was corrected for false-positive calls estimated from strand-chimera controls. Chromosomes 17–19, unused in training and model selection, formed the primary test set. In the ZF-SssI data^32^, a CG site unmethylated in all six Col-0 controls was counted as newly methylated when transgenic plants of both generations reached ≥ 0.5, and regressions controlled for neighbouring methylation, gene-body location and chromosomal region.

### CG cooperativity and the permissive background

For CG cooperativity, 200 gene-body CG sites per set-point class were held fixed while neighbouring CGs were removed or single CGs were added at defined distances, with matched non-CG substitutions as controls. To compare species, 100 of these sites per class were drawn at random and the same edited sequences were scored by all fourteen species models: removal of all other CGs within ±100 bp (CG→TG) paired with as many non-CG C→T substitutions, and one added CG (CA→CG or TG→CG) in each of five distance bands from 3 to 250 bp, each paired with a substitution of the same type that creates no CG. Effects are the edit minus its paired control at sites that the scoring model predicts to be methylable (*p* ≥ 0.3). For the permissive background, 150 gene-body CG sites in unmethylated genes (gene-mean methylation below 0.05 and *p* < 0.15) and 150 in gene-body-methylated genes (gene mean above 0.3 and *p* ≥ 0.7) on chromosome 4 were paired at random; each recipient kept its own sequence within ±*k* bp of its focal CG (*k* = 15–2,000) and took the rest of the 10-kb window from its partner, and flanks were alternatively shuffled within 1-kb blocks preserving dinucleotide or mononucleotide composition. Footprints in the measured methylomes used each species’ training methylome outside its held-out test regions: CG dyads with at least ten reads in 2-kb tiles of intermediate mean methylation (0.2–0.8), whose methylation was regressed on the numbers of other CG and of GpC dinucleotides in each distance band, with GC content as a covariate and tile fixed effects. Intervals for these analyses come from 2,000 resamples of 100-kb blocks.

For the natural-variant test, sequence variants^33^ within 3–250 bp of reference CG sites in gene bodies on chromosomes 4 and 5 were classified as creating a CG, destroying a CG or neither. Focal methylation was regressed on carrier status across 586 accessions^23^ with local background methylation and ten genotype principal components as covariates, and estimates were pooled within distance and set-point strata. The local-haplotype control compared only carriers and non-carriers with identical genotypes at all other variants within 1 kb. REED’s prediction for each pair was the change in *p* after inserting the alternative allele; the published methods were scored on the same substitutions.

### Borrowed set points

For borrowed set points, the Shaw-population (chromosome 5), Huhan3 (chromosomes 11 and 12) and poplar (chromosomes 17–19) epimutations were rescored with the frozen analysis code, replacing the target’s predictions with those of the fourteen models of other species (among the fourteen benchmark species and rice) run on the target’s own sequence; the direction of change counted as predicted when gain and loss coefficients and AUCs all had 95% intervals on the expected side, and direction AUC (the mean of the gain AUC and one minus the loss AUC) was correlated with divergence time (TimeTree 5) within each target and averaged over targets, against 100,000 within-target permutations.

### Motif grammar, DAP-seq and the CWA rule

For motifs, each of 805 JASPAR 2024 plant motifs^34^ was inserted at the centre of 10-kb backgrounds from a species’ own genome (40 moderately and 40 lowly methylated); its effect was the difference in predicted CG methylation within ±200 bp between the motif and a shuffled sequence of the same CG and CHG content; independently retrained models of seven species gave motif-effect profiles correlated at 0.81–0.95. Nucleotide attributions were gradients of predicted CG methylation with respect to the input, summarized by the bases flanking each CG. Measured methylation around native motif instances was compared with GC-matched control positions in each genome. Similarity of motif-effect profiles between species was compared with divergence times (TimeTree 5) by Mantel tests, with partial tests for model accuracy and genome-wide methylation. For per-motif calls, backgrounds were extended to 200 per species; a motif was called conserved when the combined test across species gave *q* < 0.05, at least 12 of the 14 species (including two of the three grasses) agreed in sign and the mean effect exceeded 0.5 percentage points, and one representative per JASPAR cluster was shown. Because most insertions lowered predicted methylation slightly (median −0.34 percentage points across motifs), calls were repeated after centring each species on its median motif effect; 200 motifs remained conserved. For DAP-seq^35^, binding sites of each factor detected on amplified DNA (ampDAP-seq) were labelled lost if no peak was detected on native genomic DNA (DAP-seq) within 100 bp, and REED’s predicted CG methylation within ±100 bp was compared between lost and retained sites by AUC within strata of ampDAP signal; methylation sensitivity was taken from the published motif-level DAP-seq signal in CG-methylated versus adjacent unmethylated regions. For the CWA rule, the base after CHH sites in heterochromatic transposons was switched between A and T in twelve species models (all benchmarked species except *S. viridis* and *M. esculenta*). CMT2– and RdDM-maintained regions were 200-bp tiles of chromosome 5 with CHH methylation of at least 0.05 that lost at least half of it in *cmt2* and at most a quarter in *drm1 drm2*, or the reverse^43,44^.

### Pathway resemblance

To ask which methylation pathway a model’s predictions resemble, we compared them with the methylomes of 13 *A. thaliana* pathway mutants^43^ on 200-bp segments of chromosome 4: *L* is the correlation of a model’s prediction with a mutant methylome minus its correlation with the wild-type methylome, and *D* is *L* minus the *L* of the reference *A. thaliana* model (so *D* = 0 for *A. thaliana*); for crop mutants, Δ*L* is *L* of the mutant model minus that of the wild-type model from the same study. Models trained on the 13 mutant methylomes themselves recovered the correct pathway group in every case. *E. salsugineum* lines carrying *A. thaliana CMT3*^45^ were modelled generation by generation, and tomato^46^ and rice^47^ mutant reads were realigned to their reference assemblies.

### Statistics and preregistration

Unless stated otherwise, intervals are 95% confidence intervals from 1-Mb block bootstrap (2,000 resamples), and model differences were computed on identical resamples; pathway comparisons used 100-kb blocks. Tests in the epimutation, natural-lineage, rice, poplar, SssI, natural-variant, cross-species cooperativity, methylome-footprint, borrowed-set-point, permissive-background, stress-equivalence, founder non-CG methylation and CHH-region analyses were preregistered in a version-controlled repository before the corresponding computation, and preregistration documents with their timestamps will be deposited with the code.

## Data availability

All data are public. Training methylomes: GEO GSE43857 (1001 Epigenomes), GSE75071 (GSM1942114, GSM1942115, GSM1942117, GSM1942118), GSE79526 (GSM2096945, GSM2096962, GSM2096969, GSM2096978, GSM2096980, GSM2096981, GSM2096985, GSM2096986, GSM2096987), GSM4561497 (*Z. mays*) and DDBJ DRR159267 (*O. sativa*). Epimutation data: GSE64463 (Shaw population), GSE153055 (MA3), ENA PRJEB5331 (haplogroup-1 lineage), GSE132939 (poplar; assembly GCA_024362645.1), GSE158027 (ZF-SssI), GSE223810 and GSE223861 (further accessions and biotic stress), SRA SRP047267 (soil salinity) and BioProject PRJNA319450 (rice Huhan3 and II-32B lineages). Other data: DAP-seq and ampDAP-seq peaks (GSE60143; Plant Cistrome Database), GSE39901 and GSE51304 (*A. thaliana* methylation pathway mutants), GSE128687 (*E. salsugineum* CMT3 lines) and the 1001 Genomes variant calls.

## Code availability

REED code is available at https://github.com/void-echo/REED. The trained models and genome-wide set-point maps for the fourteen benchmark species and rice, the per-site epimutation tables and the preregistration documents with their version-control timestamps will be deposited in a public repository at publication.

## Author contributions

D.W. conceived the study, developed REED, performed the analyses and wrote the manuscript. J.J. contributed to the design of the study and the interpretation of the results and revised the manuscript. J.Q. contributed to the methodology and the interpretation of the results and revised the manuscript. L.W. contributed to the conception of the study and the discussion of the results. Q.L. supervised the project and revised the manuscript. All authors read and approved the manuscript.

## Supplementary Information

**Supplementary Fig. 1.**
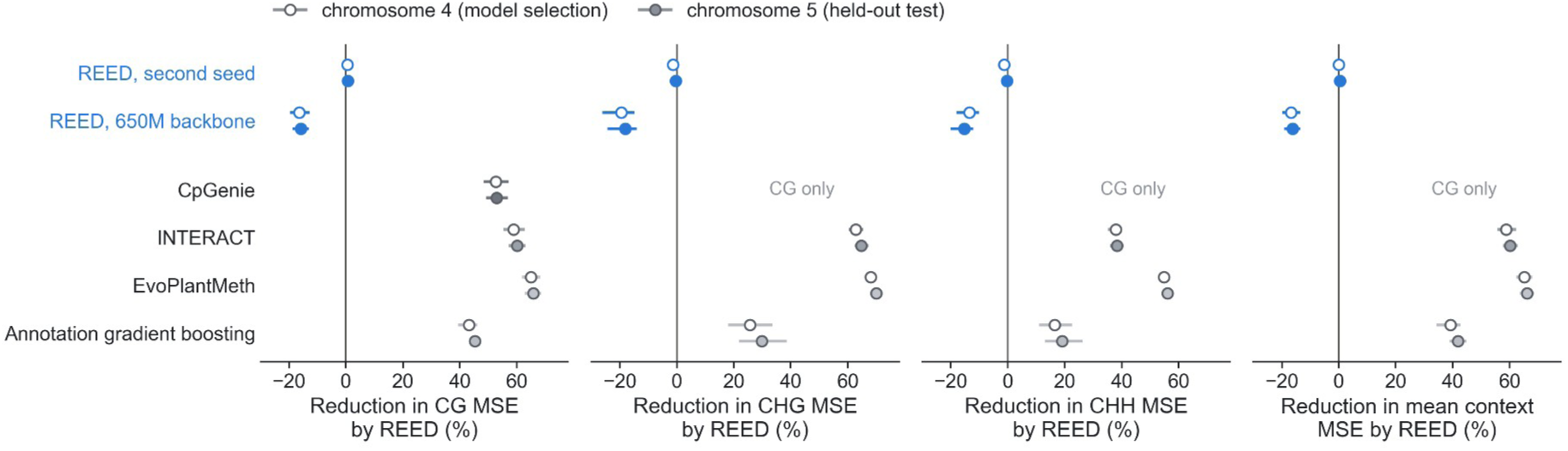
| Paired comparison on the selection and test chromosomes. Reduction in per-cytosine mean squared error of REED relative to a second REED seed, REED with the 650M-parameter backbone, three published methods and annotation gradient boosting, by context and for the mean of the three context errors, on chromosome 4 (model selection; open) and chromosome 5 (held-out test; filled); right of zero, REED has the lower error. CpGenie is compared on CG only. Bars, 95% confidence intervals from paired 1-Mb block bootstrap.

**Supplementary Fig. 2.**
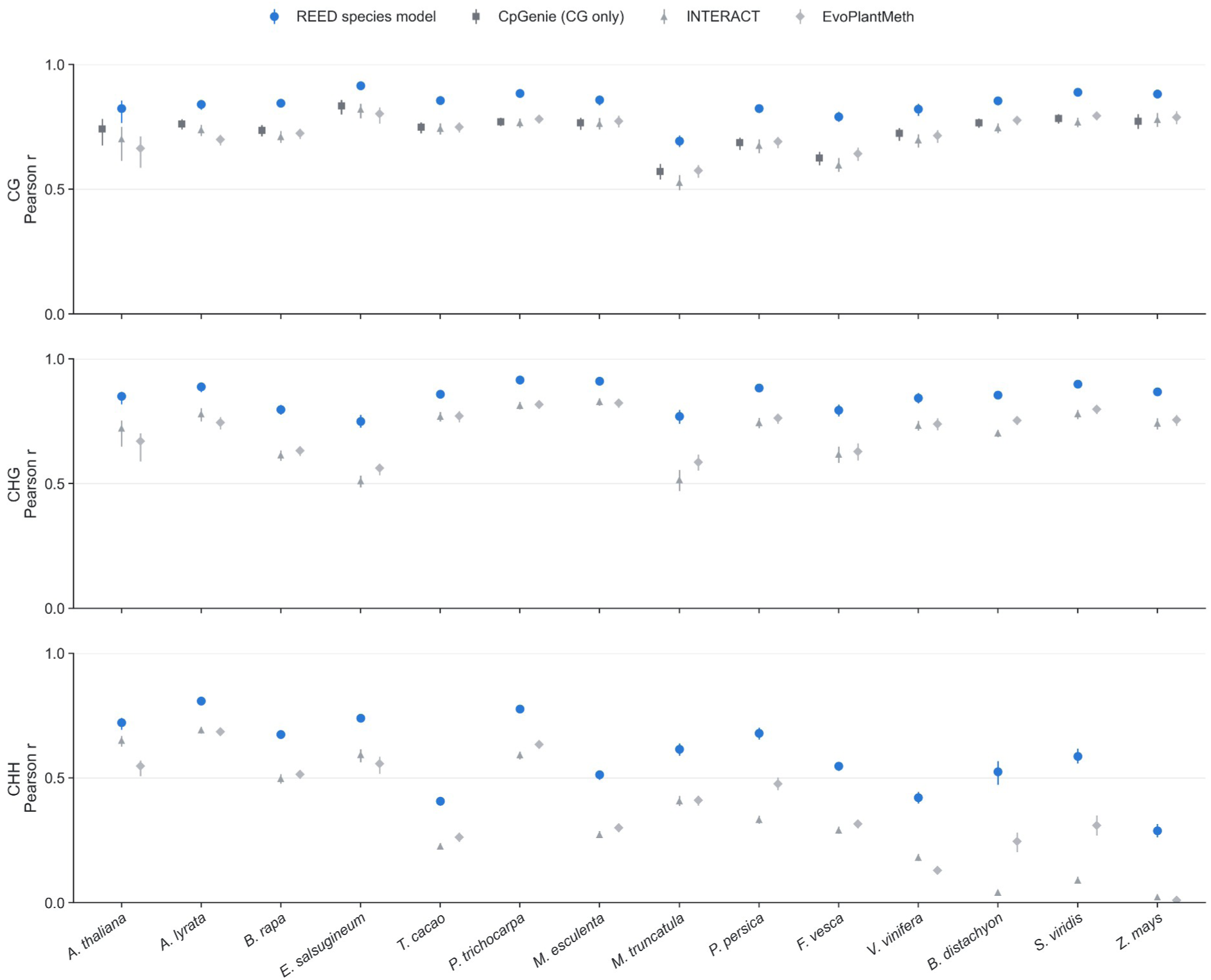
| Fourteen species, species-retrained baselines. Pearson *r* on held-out regions for REED species models and for CpGenie (CG only), INTERACT and EvoPlantMeth retrained on each species; bars, 95% intervals from 1-Mb block bootstrap.

**Supplementary Fig. 3.**
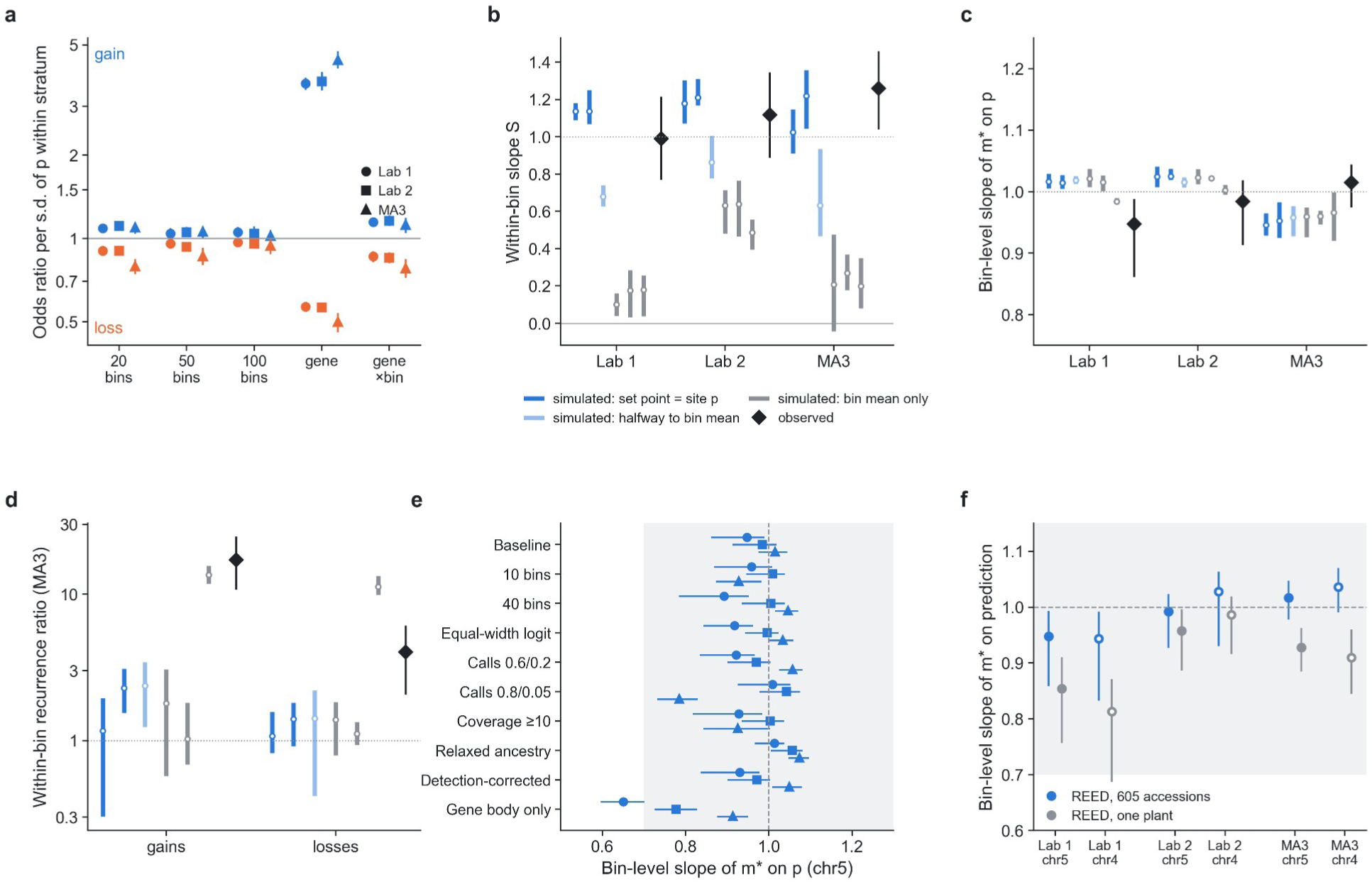
| Site-level test and robustness of the equilibrium. Within-bin and within-gene slopes *S* for the consensus and single-plant models; simulated identifiability of bin-level and within-bin statistics; equilibrium slopes, intercepts and calibration errors under alternative binning, calling thresholds, coverage, ancestral-inference rules and correction for detection asymmetry, for the Shaw population (two laboratories) and MA3 on chromosomes 4 and 5; shaded, preregistered range of the slope (0.7–1.3).

**Supplementary Fig. 4.**
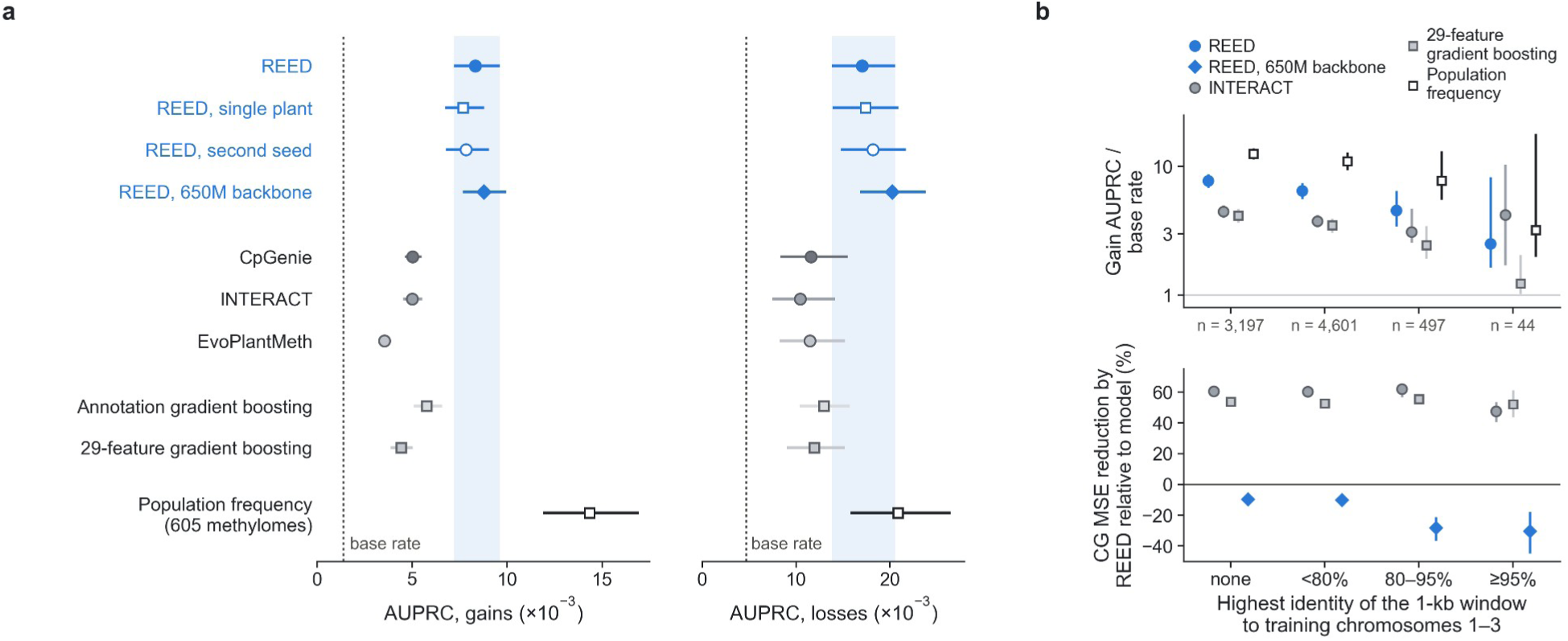
| All predictors on the same epimutations. **a**, Area under the precision–recall curve for gains (5,142 events) and losses (6,932 events) in the Shaw population (laboratory 1, chromosome 5) for REED (blue; including the model trained on a single plant, a second seed and the 650M-parameter backbone), three published methods, two gradient-boosting models and the methylation frequency of 605 accessions; dotted line, base rate; shaded band, interval of REED. **b**, Top, gain prediction (AUPRC relative to the base rate); bottom, reduction in chromosome 5 CG prediction error by REED relative to INTERACT, 29-feature gradient boosting and the 650M-parameter backbone; both stratified by the highest identity of the surrounding 1-kb window to the training chromosomes; n, gain events per stratum. Bars, 95% intervals from 1-Mb block bootstrap (paired for the error reduction).

**Supplementary Fig. 5.**
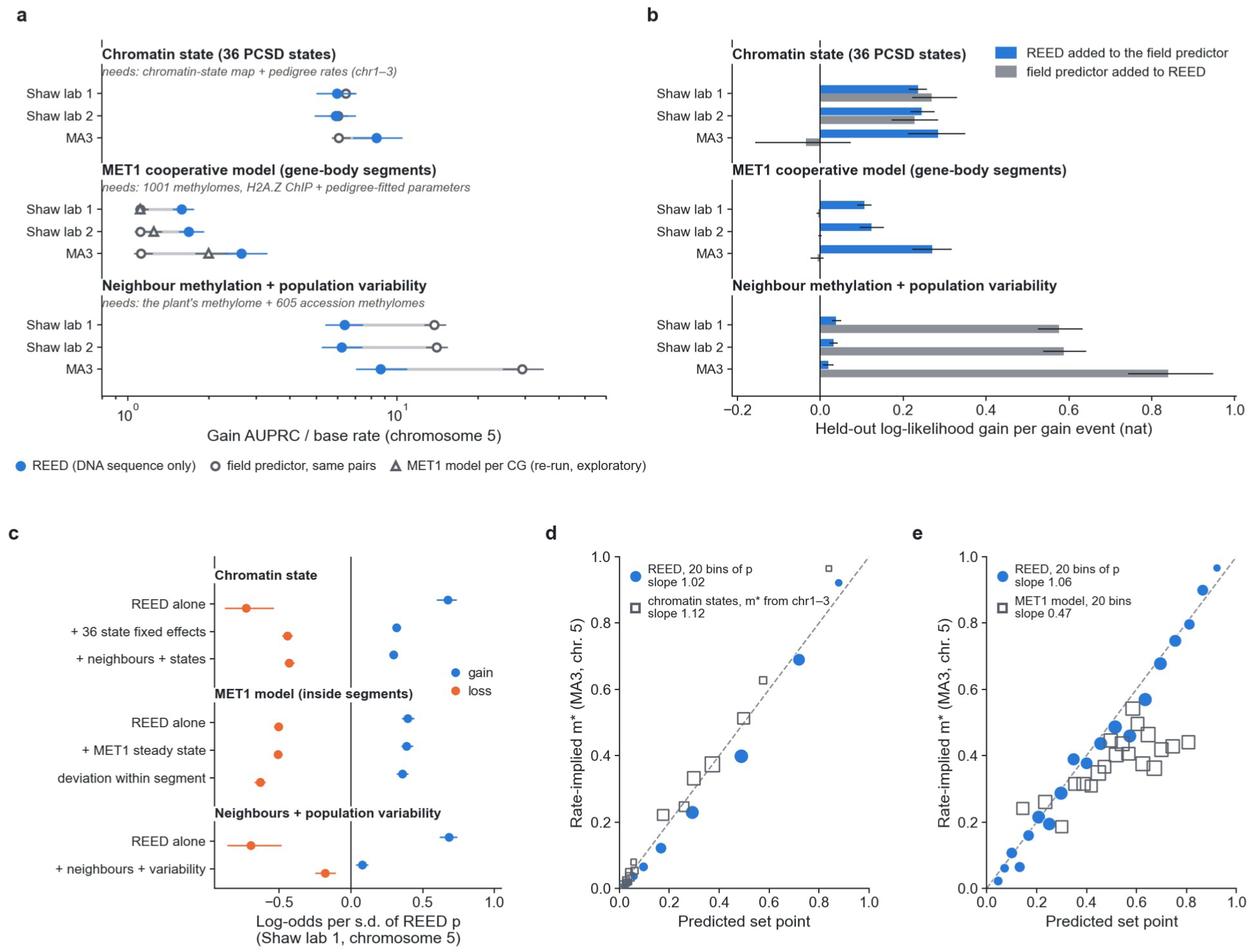
| REED and the field’s own predictors of epimutation. Each predictor is compared with REED on the sites it covers (Shaw population, laboratories 1 and 2, and MA3; chromosome 5); grey text, data each predictor requires. Chromatin states: 36 states of the Plant Chromatin State Database, with gain and loss rates estimated on chromosomes 1–3^14^. MET1 cooperative model: published steady states of gene-body segments (open circles) and, exploratory, per-CG steady states re-simulated with the published code (triangles)^11^. Neighbour methylation and population variability: neighbouring methylation of the plant and variability across 605 accessions^13^. **a**, Gain AUPRC relative to the base rate for REED (blue) and each predictor on the same pairs (open). **b**, Held-out log-likelihood gain per gain event from adding REED to the predictor (blue) or the predictor to REED (grey). **c**, Effect of REED’s *p* on gains (blue) and losses (orange) alone and after adding each predictor (Shaw population, laboratory 1). **d**, **e**, Rate-implied equilibrium against the predicted set point for REED and for chromatin states (**d**) or the MET1 model (**e**) in MA3. Bars, 95% intervals from 1-Mb block bootstrap.

**Supplementary Fig. 6.**
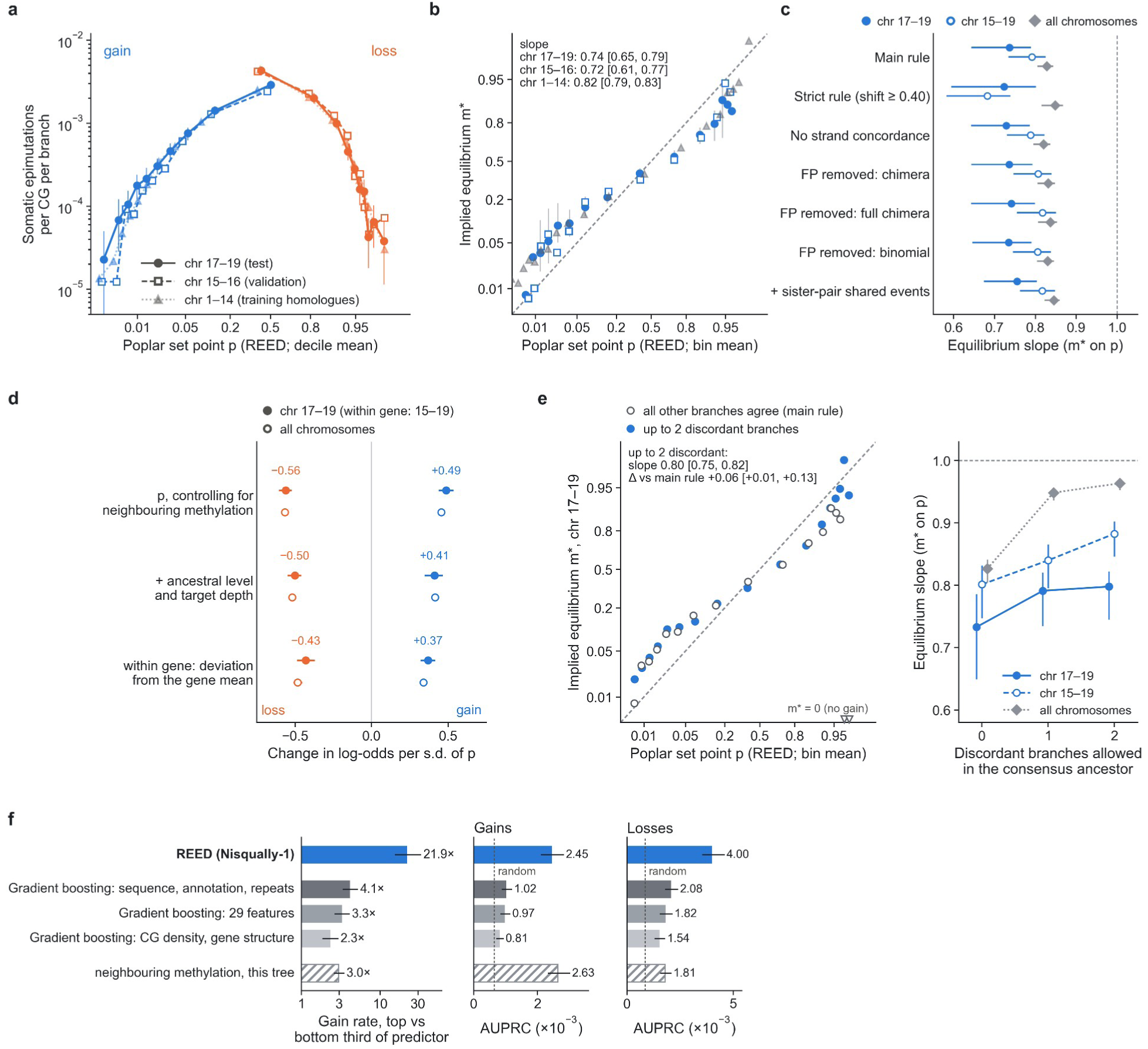
| Somatic epimutations in poplar: robustness. **a**, Single-branch somatic gain (blue; ancestrally unmethylated CGs) and loss (orange; ancestrally methylated CGs) rates per CG per branch against the poplar REED set point *p* (deciles), on chromosomes 17–19 (test; filled circles, solid), 15–16 (validation; open squares, dashed) and 1–14 (homologous to the Nisqually-1 training chromosomes; light triangles, dotted). Bars, 95% 1-Mb block-bootstrap intervals (test chromosomes). **b**, Implied equilibrium m* = gain rate / (gain + loss rate) in 20 equal-count bins of *p* for the same three chromosome groups; dashed line, m* = *p*. Weighted slopes with 95% intervals are given; intervals reaching 0 or 1 are not drawn, and bins without any gain (m* = 0) are omitted. **c**, Equilibrium slope under alternative event rules (strict: change ≥ 0.40; no requirement for strand concordance), after subtracting estimated false positives bin by bin (chimera null with real events removed; full chimera null as an upper bound; binomial sampling null) and after adding events shared by sister branches (internal-branch events), for chromosomes 17–19 (filled), 15–19 (open) and all chromosomes (grey). **d**, Coefficient of standardized *p* (log-odds per s.d.) for gain (blue) and loss (orange) in logistic models controlling for neighbouring methylation, additionally for the site’s own ancestral level and the target branch’s depth, and for the within-gene deviation of *p* from the gene mean (controlling for the gene mean and neighbouring methylation); filled, chromosomes 17–19 (within-gene model: 15–19); open, all chromosomes. **e**, Relaxing the consensus ancestor (exploratory). Left, m* against *p* on chromosomes 17–19 when every other branch must agree (main rule; open) or up to two of the other nine branches may disagree (blue); triangles at the bottom mark bins with no gain under the main rule. Right, equilibrium slope against the number of discordant branches allowed. Allowing one or two discordant branches raised the slope to 0.80 on chromosomes 17–19 and to 0.96 genome-wide, mainly by recovering gains at high-*p* sites that the strict consensus excludes. **f**, Gain enrichment (top versus bottom third of each predictor) and AUPRC for REED and for gradient-boosting models trained on the same Nisqually-1 methylome; hatched, neighbouring methylation of the old tree itself.

**Supplementary Fig. 7.**
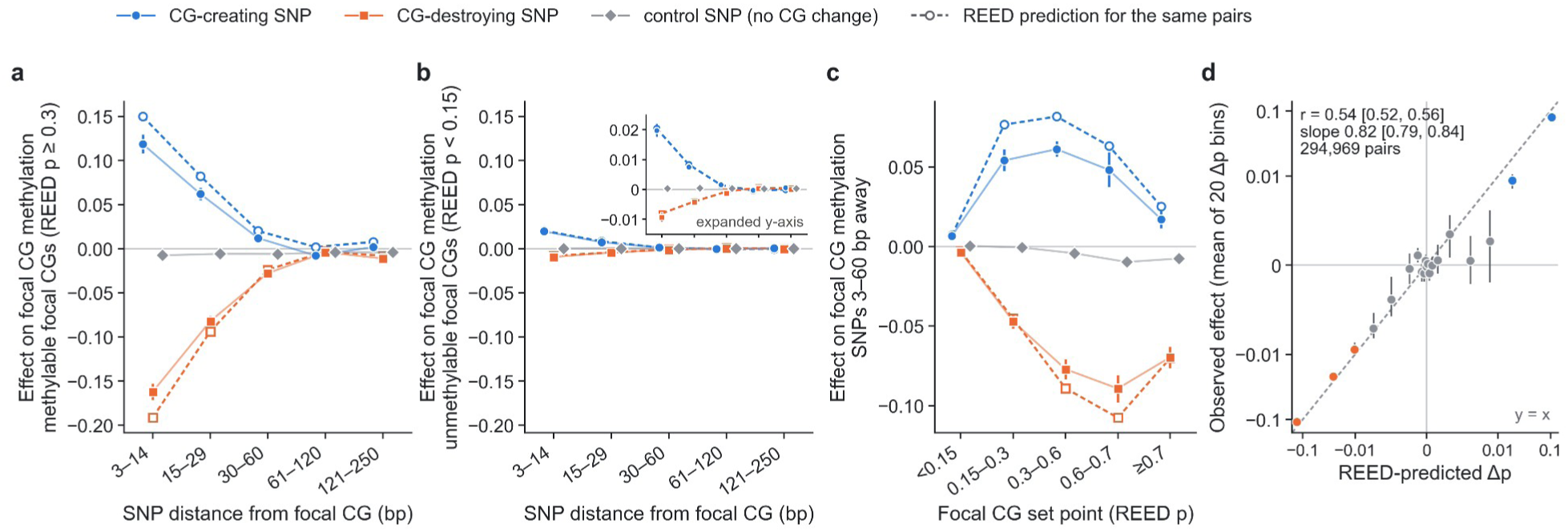
| Natural-variant test of CG cooperativity. Gene-body focal CGs on *A. thaliana* chromosomes 4 and 5, 586 accessions. Effects are the weighted mean, over variant–focal-site pairs, of the carrier coefficient in a per-pair regression of focal methylation on carrier status, local background methylation and ten genotype principal components; bars, 95% 1-Mb block-bootstrap intervals. Open symbols and dashed lines, mean REED-predicted change (Δ*p*, in silico substitution on the reference sequence) for the same pairs. **a**, **b**, Effect of SNPs that create (blue) or destroy (orange) a CG, and of control SNPs that change no CG (grey), by distance from the focal CG, at methylable (**a**; REED *p* ≥ 0.3) and low-set-point (**b**; *p* < 0.15) focal CGs; both panels share the vertical scale, and the inset in **b** shows the same data on an expanded scale. **c**, Effects of SNPs 3–60 bp from the focal CG by the focal CG’s set point. **d**, Observed effect against REED-predicted Δ*p* for all 294,969 pairs at 3–60 bp, in 20 equal-count bins of Δ*p* (both axes symmetric-log); colour, majority class in the bin (blue, CG-creating; orange, CG-destroying; grey, mixed); dashed line, y = x. Weighted r = 0.54 [0.52, 0.56]; slope 0.82 [0.79, 0.84].

**Supplementary Fig. 8.**
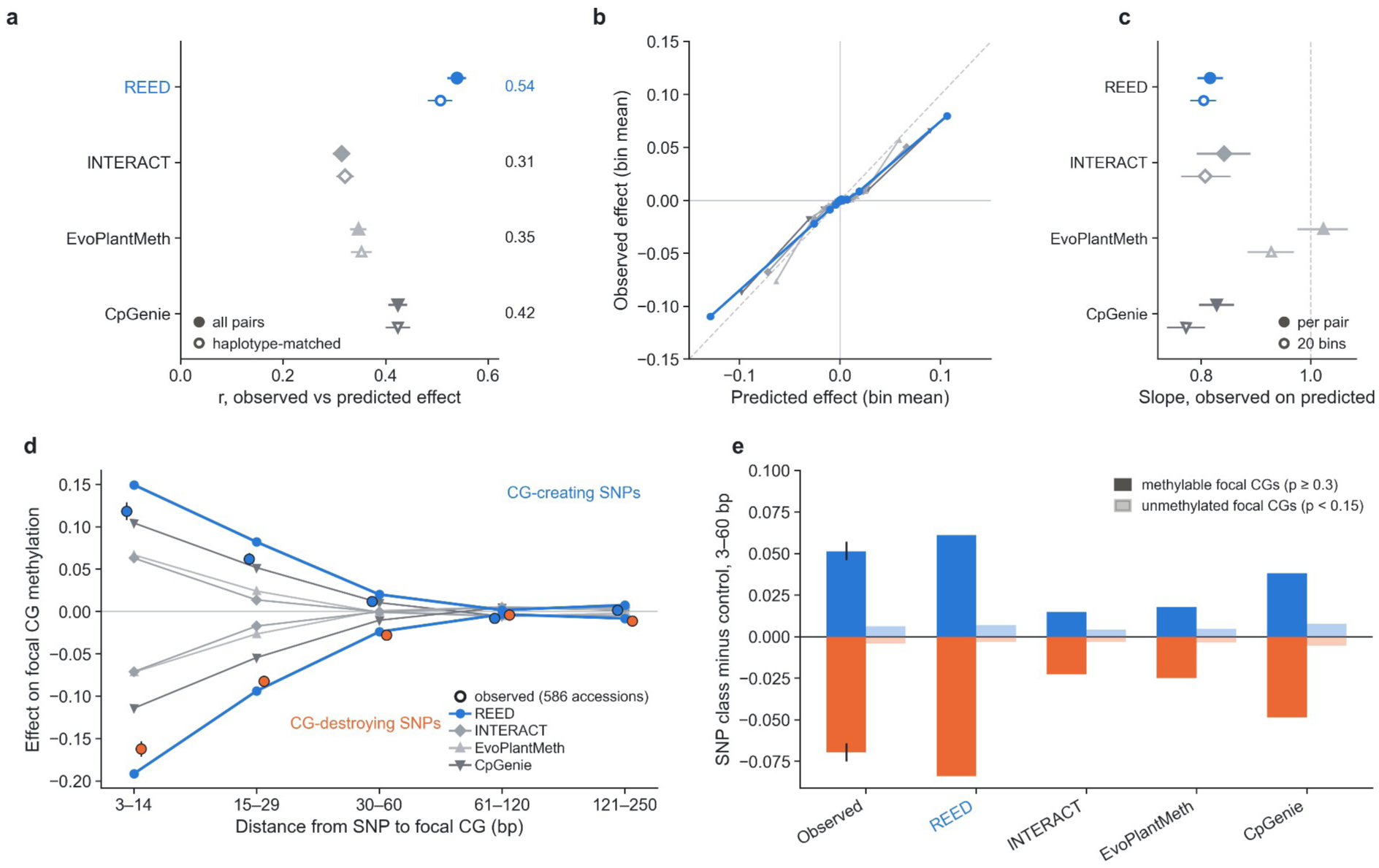
| REED and published methods on the natural-variant test. The 294,969 variant–focal-site pairs of Fig. 4b (gene-body focal CGs, variants 3–60 bp away, chromosomes 4 and 5, 586 accessions), with in silico effects computed by each model for the same substitutions. **a**, Pearson *r* between predicted and observed effects for all pairs (filled) and for the local-haplotype matched subset (open); numbers, *r* for all pairs. **b**, Observed against predicted effects in 20 bins of each model’s prediction; dashed line, identity. **c**, Slope of observed on predicted effect, per pair (filled) and over 20 bins (open). **d**, Observed effects (outlined circles) and each model’s mean prediction by distance, for CG-creating and CG-destroying variants at methylable focal CGs. **e**, Effect of CG-creating (blue) and CG-destroying (orange) variants 3–60 bp away relative to control variants, at methylable (dark) and unmethylated (light) focal CGs, observed and predicted. Error bars, 95% intervals from 1-Mb block bootstrap.

**Supplementary Fig. 9.**
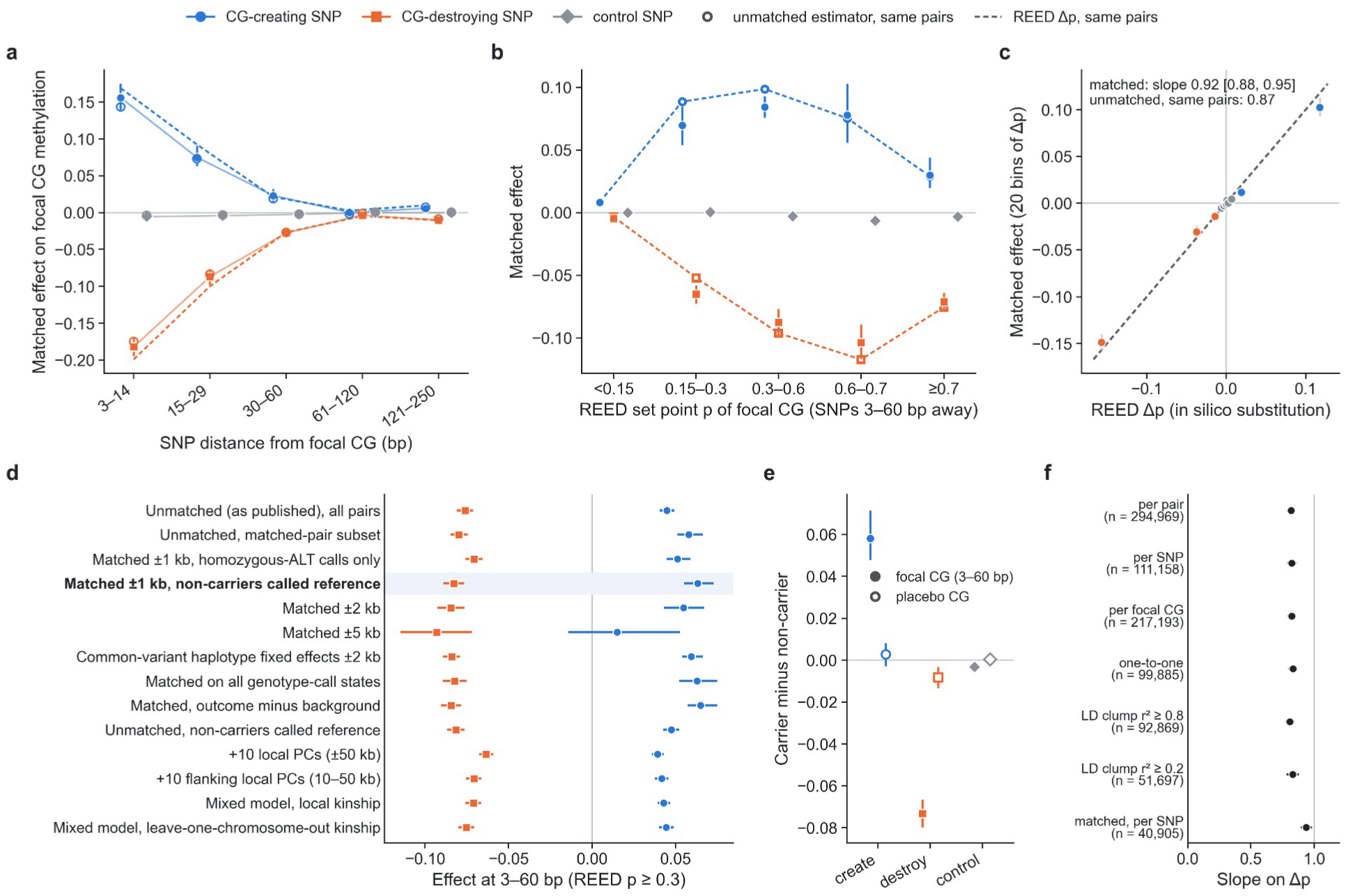
| The CG-cooperativity effect of natural variants holds within identical local haplotypes. Gene-body focal CGs on chromosomes 4 and 5, 586 accessions. Matched sets contain carriers and non-carriers with identical genotypes at all other SNPs and short indels within 1 kb of both the focal CG and the SNP; non-carriers are called homozygous reference at the tested SNP. **a**, Matched effect of CG-creating (blue), CG-destroying (orange) and control (grey) variants on focal CGs with set point *p* ≥ 0.3, by distance; open circles, unmatched estimate on the same pairs; dashed, REED Δ*p*. **b**, Matched effect at 3–60 bp by set point of the focal CG. **c**, Matched effect in 20 bins of Δ*p*; dashed line, identity. **d**, Effect at 3–60 bp under 14 estimators, including local principal components and mixed models with local or genome-wide kinship; bold, primary analysis. **e**, Difference between carriers and non-carriers in the same matched sets at the focal CG and at CGs 121–250 bp from the variant. **f**, Slope of observed on predicted effect with one estimate per pair, variant, focal CG, one-to-one pair, linkage-disequilibrium clump or matched variant. Error bars, 95% intervals from 1-Mb block bootstrap.

**Supplementary Fig. 10.**
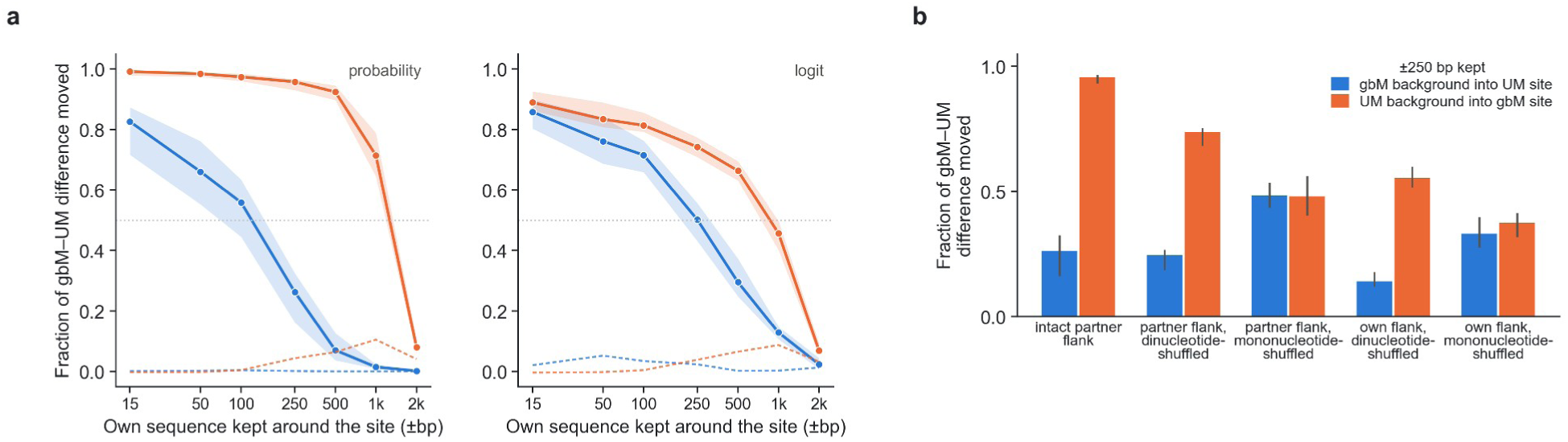
| The permissive background is read from kilobases of sequence. **a**, Background transplants between 150 pairs of gene-body CG sites in unmethylated (UM) and gene-body-methylated (gbM) *A. thaliana* genes (chromosome 4): the recipient keeps its own sequence within ±*k* bp of its focal CG and takes the rest of the 10-kb window from its partner; fraction of the gbM–UM difference in predicted methylation moved towards the partner (median over pairs; band, 95% pair-bootstrap interval) on the probability (left) and logit (right) scales; dashed, transplants between two sites of the same class. **b**, With ±250 bp kept: the intact partner flank, the partner flank shuffled preserving dinucleotide or mononucleotide composition, and the recipient’s own flank shuffled the same ways.

**Supplementary Fig. 11.**
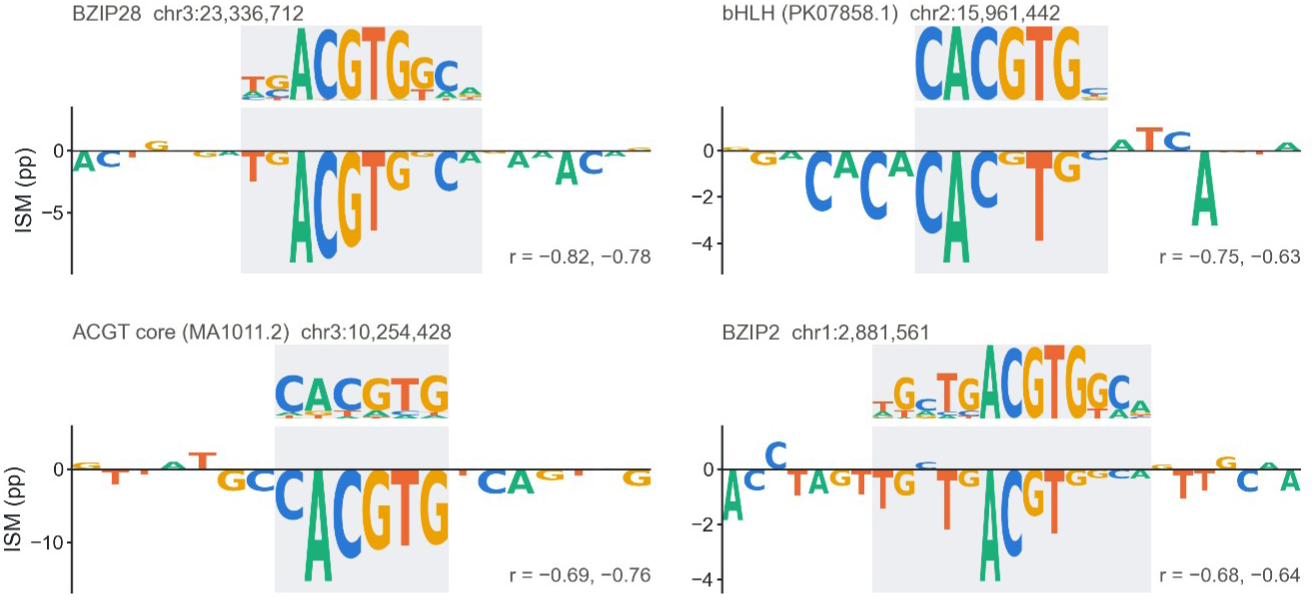
| In silico mutagenesis recovers ACGT-core motifs at genomic sites. Four *A. thaliana* sites, chosen as the strongest examples reproduced by two independently trained models (correlation between mutagenesis profile and motif ≤ −0.6 in both): JASPAR motif (top) and the effect of each reference base on predicted CG methylation of surrounding cytosines within ±200 bp (bottom; mean of two models); *r*, correlation with the motif for each model.

**Supplementary Fig. 12.**
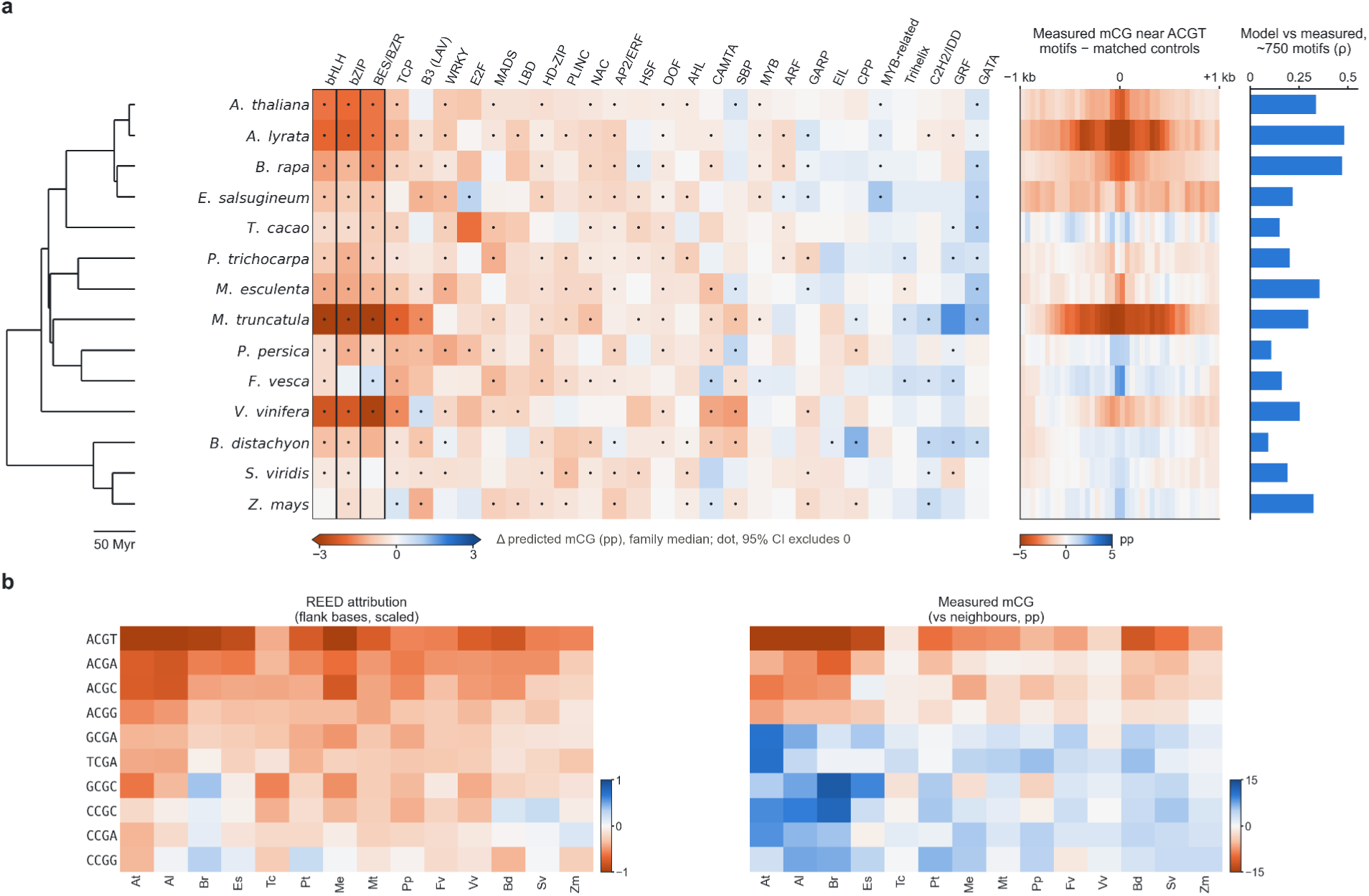
| Motif effects by transcription factor family and de novo CG-flank grammar in fourteen species. **a**, Effects by family (median over motifs; dot, 95% confidence interval excluding zero; 40 moderately methylated backgrounds per species) for fourteen species ordered by phylogeny (scale, 50 million years); right, measured CG methylation around native ACGT-core motif instances relative to matched controls in each species, and the within-species correlation between model effects and measured footprints across ∼750 motifs. **b**, De novo grammar: attribution summarized by the bases flanking each CG (left, scaled within species) and measured CG methylation of each flank combination relative to neighbouring CGs (right, centred within species).

**Supplementary Fig. 13.**
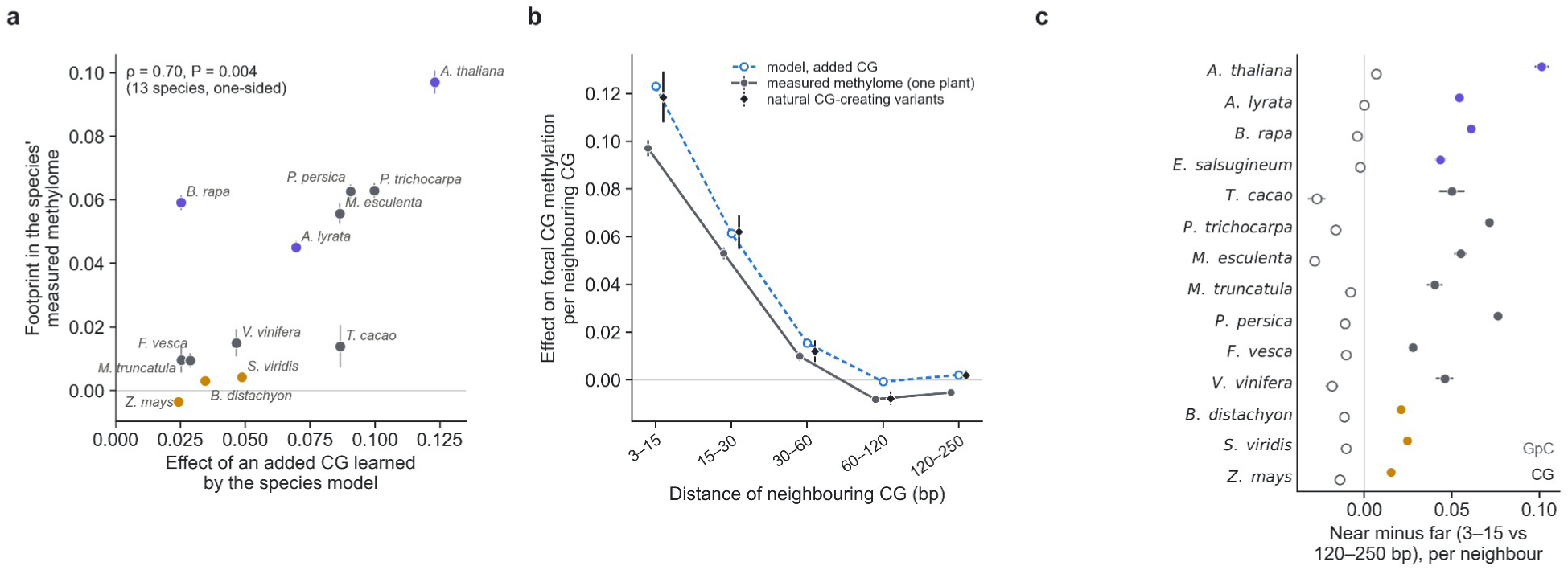
| The CG-cooperativity footprint in the measured methylomes. **a**, Effect of a CG added 3–15 bp from methylable *A. thaliana* gene-body CG sites learned by each species model (Fig. 5e) against the change in measured methylation per additional CG 3–15 bp from a focal CG in that species’ own leaf methylome (2-kb tiles of intermediate methylation, tile fixed effects, GC-adjusted); 13 species (*E. salsugineum* had too few methylable sites in the editing test); violet, Brassicaceae; dark grey, other dicots; gold, grasses. **b**, *A. thaliana*: the footprint measured in one plant’s methylome (dark grey), the model’s effect of an added CG (blue, dashed) and the effect of natural CG-creating variants among 586 accessions (black), by distance. **c**, Near-minus-far difference (3–15 bp minus 120–250 bp) per neighbouring CG (filled) and per neighbouring GpC entered in the same model (open), by species. Bars, 95% confidence intervals from 100-kb block bootstrap.

**Supplementary Fig. 14.**
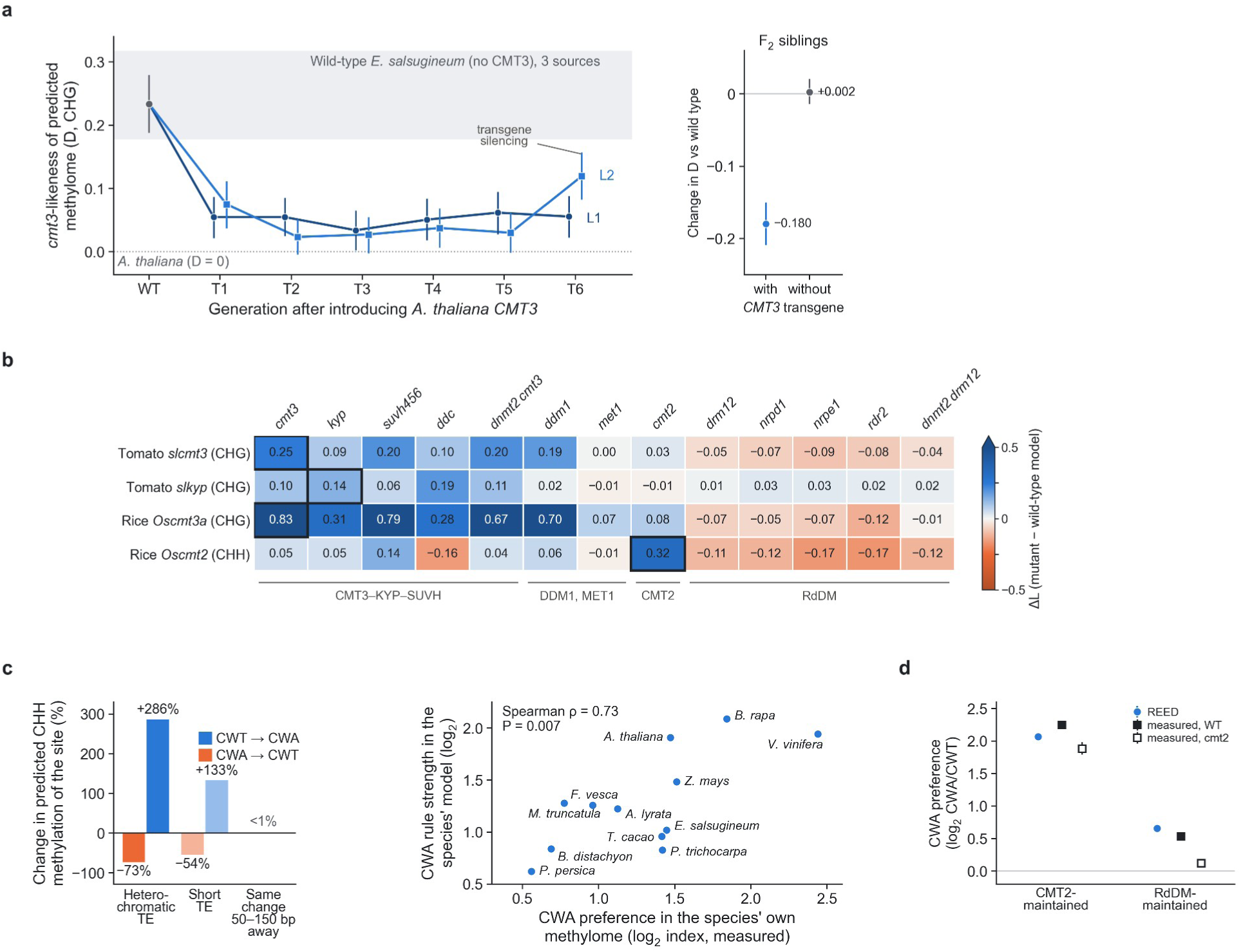
| The rules depend on the methylation enzymes each species carries. **a**, Similarity of predictions to the *A. thaliana cmt3* mutant (*D*; Methods) for models trained on wild-type *E. salsugineum* (no CMT3) and on successive generations of two lines carrying *A. thaliana CMT3*; grey band, three wild-type *E. salsugineum* methylomes; right, F2 siblings with and without the transgene. **b**, Resemblance (Δ*L*, mutant model minus same-study wild-type model) of tomato and rice mutant models to 13 *A. thaliana* pathway mutants^43^; outlined, the expected mutant. **c**, The CHH rule of the chromomethylase CMT2: changing CWA to CWT at a CHH site lowers its predicted methylation by 73% in heterochromatic transposons (reverse change, +286%; lighter bars, short transposons; right, the same change 50–150 bp away), and across twelve species models the strength of this rule tracks each species’ measured CWA preference. **d**, The same rule in CHH regions maintained by CMT2 or by RNA-directed DNA methylation (tiles defined from *cmt2* and *drm1 drm2* methylomes, chromosome 5): REED’s rule strength (log2 of predicted CWA over CWT methylation after switching the base) and the CWA/CWT index measured in wild-type and *cmt2* methylomes. Error bars in **a**, **b** and **d**, 95% confidence intervals (100-kb block bootstrap).

